# Onset, timing, and exposure therapy of stress disorders: mechanistic insight from a mathematical model of oscillating neuroendocrine dynamics

**DOI:** 10.1101/042887

**Authors:** Lae Kim, Maria D’Orsogna, Tom Chou

**Affiliations:** Dept. of Biomathematics, Univ of California, Los Angeles, Los Angeles, USA.; Department of Mathematics, CalState-Northridge, Los Angeles, USA.

**Keywords:** HPA-axis, PTSD, Stress Disorders, Dynamical system

## Abstract

The hypothalamic-pituitary-adrenal (HPA) axis is a neuroendocrine system that regulates numerous physiological processes. Disruptions in the activity of the HPA axis are correlated with many stress-related diseases such as post-traumatic stress disorder (PTSD) and major depressive disorder. In this paper, we characterize "normal” and "diseased” states of the HPA axis as basins of attraction of a dynamical system describing the inhibition of peptide hormones such as corticotropin-releasing hormone (CRH) and adrenocorticotropic hormone (ACTH) by circulating glucocorticoids such as cortisol (CORT). In addition to including key physiological features such as ultradian oscillations in cortisol levels and self-upregulation of CRH neuron activity our model distinguishes the relatively slow process of cortisol-mediated CRH biosynthesis from rapid trans-synaptic effects that regulate the CRH secretion process. Crucially we find that the slow regulation mechanism mediates external stress-driven transitions between the stable states in novel, intensity, duration, and timing-dependent ways. These results indicate that the *timing* of traumatic events may be an important factor in determining if and how patients will exhibit hallmarks of stress disorders. Our model also suggests a mechanism whereby exposure therapy of stress disorders such as PTSD may act to normalize downstream dysregulation of the HPA axis.

## Introduction

Stress is an essential component of an organism’s attempt to adjust its internal state in response to environmental change. The experience, or even the perception of physical and/or environmental change, induces stress responses such as the secretion of glucocorticoids hormones (CORT) – cortisol in humans and corticosterone in rodents – by the adrenal gland. The adrenal gland is one component of the hypothalamic-pituitary-adrenal (HPA) axis, a collection of interacting neuroendocrine cells and endocrine glands that play a central role in stress response. The basic interactions involving the HPA axis are shown in Fig. 1.. The paraventricular nucleus (PVN) of the hypothalamus receives synaptic inputs from various neural pathways via the central nervous system that are activated by both cognitive and physical stressors. Once stimulated, CRH neurons in the PVN secrete corticotropin-releasing hormone (CRH), which then stimulates the anterior pituitary gland to release adrenocorticotropin hormone (ACTH) into the bloodstream. ACTH then activates a complex signaling cascade in the adrenal cortex, which ultimately releases glucocorticoids (Fig. 1B). In return, glucocorticoids exert a negative feedback on the hypothalamus and pituitary, suppressing CRH and ACTH release and synthesis in an effort to return them to baseline levels. Classic stress responses include transient increases in levels of CRH, ACTH, and cortisol. The basic components and organization of the vertebrate neuroendocrine stress axis arose early in evolution and the HPA axis, in particular, has been conserved across mammals [1].

**Figure 1:**
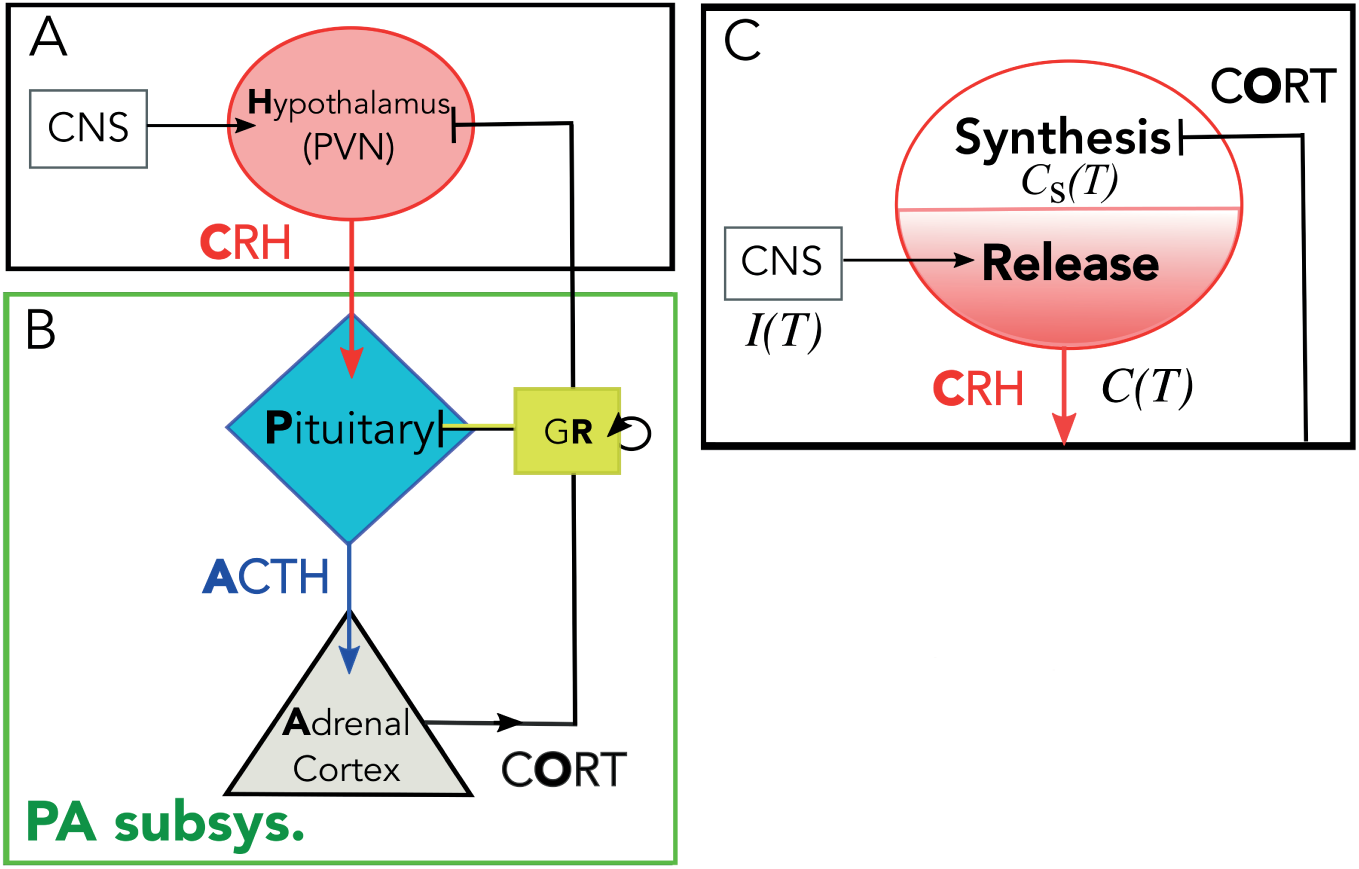
Schematic of HPA axis. (A) Stress is processed in the central nervous system (CNS) and a signal is relayed to the PVN in the hypothalamus to activate CRH secretion into the hypophyseal portal system. (B) CRH diffuses to the pituitary gland and activate ACTH secretion. ACTH travels down to the adrenal cortex to activate cortisol (CORT) release. Cortisol inhibits both CRH and ACTH secretion to down-regulate its own production, forming a closed loop. In the pituitary gland, cortisol binds to glucocorticoid receptors (GR) (yellow box) to inhibit ACTH and self-upregulate GR production. This part of the axis comprises the PA subsystem. (C) Negative feedback of cortisol affects the synthesis process in the hypothalamus, which indirectly suppresses the release of CRH. External inputs such as stressors and circadian inputs directly affect the release rate of the CRH.

Dysregulation in the HPA axis is known to correlate with a number of stress-related disorders. Increased cortisol (hypercortisolism) is associated with major depressive disorder (MDD) [2, 3], while decreased cortisol (hypocortisolism) is a feature of post-traumatic stress disorder (PTSD), post infectious fatigue, and chronic fatigue syndrome (CFS) [4–7]. Since PTSD develops in the aftermath of extreme levels of stress experienced during traumatic incidents like combat, sexual abuse, or life-threatening accidents, its progression may be strongly correlated with disruption of the HPA axis caused by stress response. For example, lower peak and nadir cortisol levels were found in patients with combat-related PTSD [8].

Mathematical models of the HPA axis have been previously formulated in terms of dynamical systems of ordinary differential equations (ODEs) [9–12] or delay differential equations (DDEs) [13–15] that describe the time-evolution of the key regulating hormones of the HPA axis: CRH, ACTH, and cortisol. These models [13, 14, 16] incorporate positive self-regulation of glucocorticoid receptor expression in the pituitary, which may generate bistability in the dynamical structure of the model [17]. Of the two stable equilibrium states, one is characterized by higher levels of cortisol and is identified as the “normal” state. The other is characterized by lower levels of cortisol and can be interpreted as one of the “diseased” states associated with *hypocortisolism.* Stresses that affect the activity of neurons in the PVN are described as perturbations to endogenous CRH secretion activity. Depending on the length and magnitude of the stress input, the system may or may not shift from the basin of attraction of the normal steady state towards that of the diseased one. If such a transition does occur, it may be interpreted as the onset of disease. A later model [16] describes the effect of stress on the HPA axis as a gradual change in the parameter values representing the maximum rate of CRH production and the strength of the negative feedback activity of cortisol. Changes in cortisol secretion pattern are assumed to arise from anatomical changes that are mathematically represented as changes to the corresponding parameter values [16].

Both classes of models imply qualitatively different time courses of disease progression [16, 17]. The former suggests that the abnormal state is a pre-existing basin of attraction of a dynamical model that stays dormant until a sudden transition is triggered by exposure to trauma [17]. In contrast, the latter assumes that the abnormal state is reached by the slow development of structural changes in physiology due to the traumatic experience [16]. Although both models [16, 17] describe changes in hormonal levels experienced by PTSD patients, they both fail to exhibit stable ultradian oscillations in cortisol, which is known to play a role in determining the responsiveness of the HPA axis to stressors [18].

In this study, we consider a number of distinctive physiological features of the HPA axis that give a more complete picture of the dynamics of stress disorders and that have not been considered in previous mathematical models. These include the effects of intrinsic ultradian oscillations on HPA dysregulation, distinct rapid and slow feedback actions of cortisol, and the correlation between HPA imbalance and disorders induced by external stress. As with the majority of hormones released by the body, cortisol levels undergo a circadian rhythm, starting low during night sleep, rapidly rising and reaching its peak in the early morning, then gradually falling throughout the day. Superposed on this slow diurnal cycle is an ultradian rhythm consisting of approximately hourly pulses. CRH, ACTH, and cortisol are all secreted episodically, with the pulses of ACTH slightly preceding those of cortisol [19].

As for many other hormones such as gonadotropin-releasing hormone (GnRH), insulin, and growth hormone (GH), the ultradian release pattern of glucocorticoids is important in sustaining normal physiological functions, such as regulating gene expression in the hippocampus [20]. It is unclear what role oscillations play in homeostasis, but the time of onset of a stressor in relation to the phase of the ultradian oscillation has been shown to influence the physiological response elicited by the stressor [21].

To distinguish the rapid and slow actions of cortisol, we separate the dynamics of biosynthesis of CRH from its secretion process, which operate over very different timescales [22]. While the two processes are mostly independent from each other, the rate of CRH secretion should depend on the synthesis process since CRH peptides must be synthesized first before being released (Fig. 1C). On the other hand, the rate of CRH peptide synthesis is influenced by cortisol levels, which in turn, are regulated by released CRH levels. We will investigate how the separation and coupling of these two processes can allow stress-induced dysregulations of the HPA axis.

The mathematical model we derive incorporates the above physiological features and reflects the basic physiology of the HPA axis associated with delays in signaling, fast and slow negative feedback mechanisms, and CRH self-upregulation [23]. Within an appropriate parameter regime, our model exhibits two distinct stable *oscillating* states, of which one is marked by a larger oscillation amplitude and a higher base cortisol level than the other. These two states will be referred to as normal and diseased states. Our interpretation is reminiscent of the two-state dynamical structure that arises in the classic Fitzhugh-Nagumo model of a single neuron, in which resting and spiking states emerge as bistable modes of the model [24], or in models of neuronal networks where an “epileptic brain” is described in terms of the distance between a normal and a seizure attractor in phase-space [25].

## Models

Models of HPA dynamics [13, 14, 16, 17, 26] are typically expressed in terms of ordinary differential equations (ODEs):

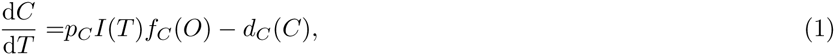

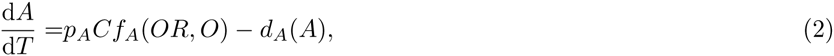

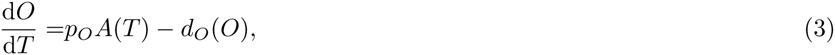

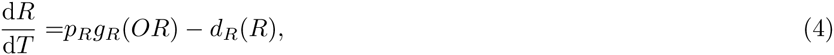

where *C*(*T*), *A*(*T*), and *O*(*T*) denote the plasma concentrations of CRH, ACTH, and cortisol at time *T*, respectively. *R*(*T*) represents the availability of glucocorticoid receptor (GR) in the anterior pituitary. The amount of cortisol bound GR is typically in quasi-equilibrium so concentration of the ligand-receptor complex is approximately proportional to the product *O*(*T*)*R*(*T*) [17]. The parameters *p*_*α*_ (*α* ∈ {*C*,*A*, *O*, *R*}) relate the production rate of each species *α* to specific factors that regulate the rate of release/synthesis. External stresses that drive CRH release by the PVN in the hypothalamus are represented by the input signal *I*(*T*). The function *f*_*C*_(*O*) describes the negative feedback of cortisol on CRH levels in the PVN while *f*_*A*_(*OR*, *O*) describes the negative feedback of cortisol or cortisol-GR complex (at concentration *O*(*T*)*R*(*T*)) in the pituitary. Both are mathematically characterized as being positive, decreasing functions so that *f*_*A*,*C*_(·) ≥ 0 and 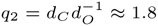. On the other hand, the function *g*_*R*_(*OR*) describes the self-upregulation effect of the cortisol-GR complex on GR production in the anterior pituitary [27]. In contrast to *f*_*A*,*C*_(·), *g*_*R*_(·) is a positive but increasing function of *OR* so that *g*_*R*_(·) ≥ 0 and 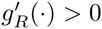. Finally, the degradation functions *d*_*α*_(*·*) describe how each hormone and receptor is cleared and may be linear or nonlinear.

Without including the effects of the glucocorticoid receptor (neglecting Eq. 4 and assuming *f*_*A*_(*OR*, *O*) = *f*_*A*_(*O*) in Eq. 2), Eqs. 1-3 form a rudimentary “minimal” model of the HPA axis [9, 28]. If *f*_*A*, *C*_(·) are Hill-type feedback functions dependent only on *O*(*T*) and *d*_*α*_(·) are linear, a unique global stable point exists. This equilibrium point transitions to a limit cycle through a Hopf bifurcation but only within nonphysiological parameter regimes [9]. The inclusion of GR and its self-upregulation in the anterior pituitary [17] creates two stable equilibrium states of the system, but still does not generate oscillatory behavior. More recent studies extend the model (represented by Eq. 1-4) to include nonlinear degradation [16] or constant delay to account for delivery of ACTH and synthesis of glucocorticoid in the adrenal gland [13]. These two extended models exhibit only one intrinsic circadian [16] or ultradian [13] oscillating cycle for any given set of parameter values, precluding the interpretation of normal and diseased states as bistable oscillating modes of the model.

Here, we develop a new model of the HPA axis by first adapting previous work [13] where a physiologically-motivated delay was introduced into Eq. 3, giving rise to the observed ultradian oscillations [13]. We then improve the model by distinguishing the relatively slow mechanism underlying the cortisol-mediated CRH biosynthesis from the rapid trans-synaptic effects that regulate the CRH secretion process. This allows us to decompose the dynamics into slow and fast components. Finally, self-upregulation of CRH release is introduced which allows for bistability. These ingredients can be realistically combined in a way that leads to novel, clinically identifiable features and are systematically developed below

### Ultradian rhythm and time delay

Experiments on rats show a 3-6 minute inherent delay in the response of the adrenal gland to ACTH [29]. Moreover, in experiments performed on sheep [30], persistent ultradian oscillations were observed even after surgically removing the hypothalamus, implying that oscillations are inherent to the PA subsystem. Since oscillations can be induced by delays, we assume, as in Walker *et al.* [13], a time delay *T*_d_ in the ACTH-mediated activation of cortisol production downstream of the hypothalamus. Eq. 3 is thus modified to

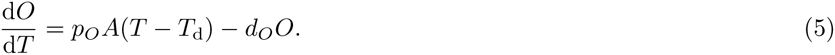

Walker *et al*. [13] show that for fixed physiological levels of CRH, the solution to Eqs. 2, 4 and 5 leads to oscillatory *A*(*T*), *O*(*T*), and *R*(*T*). In order to describe the observed periodic cortisol levels in normal and diseased states, the model requires *two* oscillating stable states. We will see that dual oscillating states arise within our model when the delay in ACTH-mediated activation of cortisol production is coupled with other known physiological processes.

### Synthesis of CRH

CRH synthesis involves various pathways, including CRH gene transcription and transport of packaged CRH from the cell body (soma) to their axonal terminals where they are stored prior to release. Changes in the steady state of the synthesis process typically occur on a timescale of minutes to hours. On the other hand, the secretory release process depends on changes in membrane potential at the axonal terminal of CRH neurons, which occur over millisecond to second timescales.

To model the synthesis and release process separately, we distinguish two compartments of CRH: the concentration of stored CRH within CRH neurons will be denoted *C*_s_(*T*), while levels of released CRH in the portal vein outside the neurons will be labeled *C*(*T*) (Fig. 1.C). Newly synthesized CRH will first be stored, thus contributing to *C*_s_. We assume that the stored CRH level *C*_s_ relaxes toward a target value set by the function *C*_∞_(*O*):

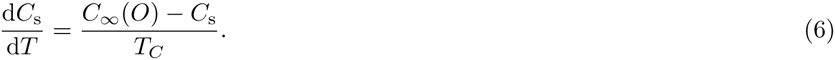

Here, *T*_*C*_ is a characteristic time constant and *C*_∞_(*O*) is the *Cortisol-dependent* target level of stored CRH. Eq. 6 also assumes that the relatively small amounts of CRH released into the bloodstream do not significantly deplete the *C*_s_ pool. Note that the effects induced by changing cortisol levels are immediate as the production term *C*_∞_(*O*)/*T*_*C*_ is adjusted instantaneously to current cortisol levels. Our model thus does not exclude cortisol rapidly acting on the initial transcription activity, as suggested by CRH hnRNA (precursor mRNA) measurements [31]. On the other hand, the time required to reach the steady state for the completely synthesized CRH peptide will depend on the characteristic time scale constant *T*_*C*_. Ideally, *T*_*C*_ should be estimated from measurements of the pool size of releasable CRH at the axonal terminals. To best of our knowledge, there are currently no such measurements available, so we base our estimation on mRNA level measurements. We believe this is a better representation of releasable CRH than hnRNA levels since mRNA synthesis is a further downstream process. Previous studies have shown that variations in CRH mRNA due to changes in cortisol levels take at least twelve hours to detect [32]. Therefore, we estimate *T*_*C*_ ≳ 12hrs = 720min. The negative feedback of cortisol on CRH levels thus acts through the production function *C*_∞_(*O*) on the relatively slow timescale *T*_C_. To motivate the functional form of *C*_∞_(*O*), we invoke experiments on rats whose adrenal glands had been surgically removed and in which glucocorticoid levels were subsequently kept fixed (by injecting exogenous glucocorticoid) for 5-7 days [22, 33]. The measured CRH mRNA levels in the PVN were found to decrease exponentially with the level of administered glucocorticoid [22, 33]. Assuming the amount of releasable CRH is proportional to the amount of measured intracellular CRH mRNA, we can approximate *C*_∞_(*O*) as a decreasing exponential function of cortisol level *O*.

### Secretion of CRH

To describe the CRH secretion, we consider the following three factors: synaptic inputs to CRH cells in the PVN, availability of releasable CRH peptide, and self-upregulation of CRH release.

CRH secretion activity is regulated by synaptic inputs received by the PVN from multiple brain regions including limbic structures like the hippocampus and the amygdala, that are activated during stress. It has been reported that for certain types of stressors, these synaptic inputs are modulated by cortisol independent of, or parallel to, its regulatory function on CRH synthesis activity [34]. On the other hand, a series of studies [35–37] showed that cortisol did not affect the basal spiking activity of the PVN. We model the overall synaptic input, denoted by *I*(*T*) in Eq. 1, as follows

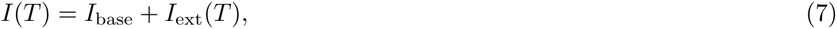

where *I*_base_ and *I*_ext_(*t*) represent the basal firing rate and stress-dependent synaptic input of the PVN, respectively. As the effect of cortisol on the synaptic input during stress is specific to type of stressor [38–40], we assume *I*_ext_(*T*) to be independent of *O* for simplicity and generality. Possible implications of cortisol dependent input function *I*_ext_(*T*, *O*) on model behavior will be discussed in the Additional File.

The secretion of CRH will also depend upon the amount of stored *releasable* CRH, *C*_s_(*T*), within the neuron and inside the synaptic vesicles. Therefore, *C*_s_ can also be factored into Eq. 1 through a source term *h*(*C*_s_) which describes the amount of CRH released per unit of action potential activity of CRH neurons. Finally, it has been hypothesized that CRH enhances its own release [23], especially when external stressors are present. The enhancement of CRH release by CRH is mediated by activation of the membrane-bound G-protein-coupled receptor CRHR-1 whose downstream signaling pathways operate on timescales from milliseconds to seconds [41, 42]. Thus, self-upregulation of CRH release can be modeled by including a positive and increasing function *g*_*C*_ (*C*) in the source term in Eq. 1.

Combining all these factors involved in regulating the secretion process, we can rewrite Eq. 1 by replacing *f*_*C*_(*O*) with *h*(*C*_s_)*g*_*C*_(*C*) as follows

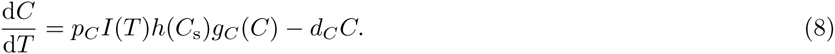

In this model (represented by Eqs. 6,8,2,5, and 4), cortisol no longer *directly* suppresses CRH levels, rather, it decreases CRH synthesis through Eq. 6, in turn suppressing *C*_s_. The combination *h*(*C*_s_)*g*_*C*_(*C*) in Eq. 8 indicates the release rate of stored CRH decreases when either *C*_s_ or *C* decreases. We assume that inputs into the CRH neurons modulate the overall release process with weight *p*_*C*_.

### Complete delay-differential equation model

We are now ready to incorporate the mechanisms described above into a new, more comprehensive mathematical model of the HPA axis, which, in summary, includes

i. A delayed response of the adrenal cortex to cortisol (Eq. 5).
ii. A slow time-scale negative feedback by cortisol on CRH synthesis (through the *C*_∞_(*O*) production term in Eq. 6).
iii. A fast-acting positive feedback of stored and circulating CRH on CRH release (through the *h*(*C*_s_)*g*_*C*_(*C*) term in Eq. 8);

Our complete mathematical model thus consists of Eqs. 2, 4, 5, 6, and 8. We henceforth assume *f*_*A*_(*OR*,*O*) = *f*_*A*_(*OR*) depends on only the cortisol-GR complex and use Hill-type functions for *f*_*A*_(*OR*) and *g*_*R*_(*OR*) [13, 14, 16, 17]. Our full theory is characterized by the following system of delay differential equations:

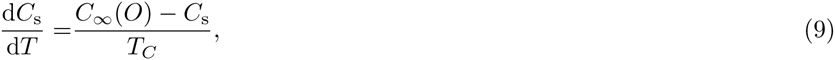

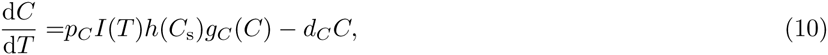

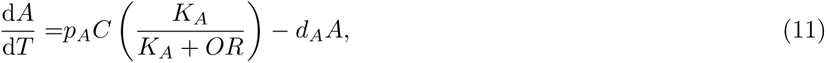

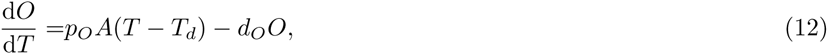

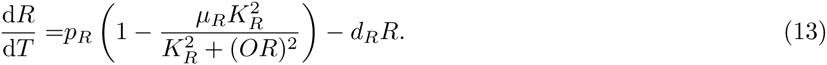

The parameters *K*_*A*, *R*_ represent the level of *A* and *R* at which the negative or positive effect are at their half maximum and 1 – *μ*_*R*_ represents the basal production rate for GR when *OR* = 0.

Of all the processes modeled, we will see that the slow negative feedback will be crucial in mediating transitions between stable states of the system. The slow dynamics will allow state variables to cross basins of attraction associated with each of the stable states.

### Nondimensionalization

To simplify the further development and analysis of our model, we nondimensionalize Eqs. 9-13 by rescaling all variables and parameters in a manner similar to that of Walker *et al*. [13], as explicitly shown in the Additional File. We find

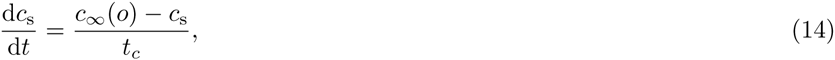

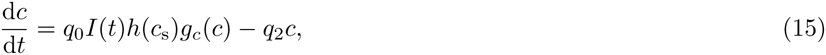

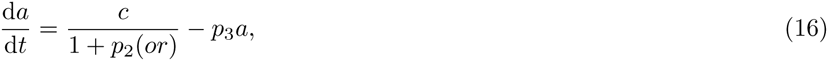

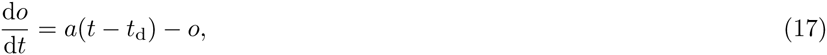

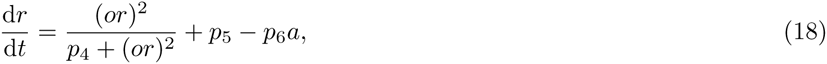

where *c*_*s*_, *c*, *a*, *r*, *o* are the dimensionless versions of the original concentrations *C*_*s*_, *C*, *A*, *R*, *O*, respectively. The dimensionless delay in activation of cortisol production by ACTH is now denoted *t*_d_. All dimensionless parameters *q*_*i*_, *p*_*i*_, *t*_d_, and *t*_c_ are combinations of the physical parameters and are explicitly given in the Additional File. The functions *c*_∞_(*o*), *h*(*c*_s_), and *g*_*c*_(*c*) are dimensionless versions of *C*_∞_(*O*), *h*(*C*_s_), and *g*_*C*_(*C*), respectively, and will be chosen phenomenologically to be

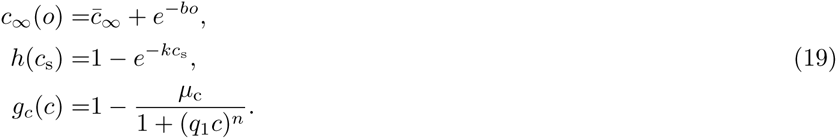

The form of *c*_∞_(*o*) is based on the above-mentioned exponential relation observed in adrenalectomized rats [22, 33]. The parameters *C̄*_∞_ and *b* represent the minimum dimensionless level of stored CRH and the decay rate of the function, respectively. How the rate of CRH release increases with *c*_s_ is given by the function *h*(*c*_s_). Since the amount of CRH packaged in release vesicles is likely regulated, we assume *h*(*c*_s_) saturates at high *c*_s_. The choice of a decreasing form for *c*_∞_(*o*) implies that increasing cortisol levels will decrease the target level (or production rate) of *c*_s_ in Eq. 14. The reduced production of *c*_s_ will then lead to a smaller *h*(*c*_s_) and ultimately a reduced release source for *c* (Eq. 15). As expected, the overall effect of increasing cortisol is a decrease in the release rate of CRH. Finally, since the upregulation of CRH release by circulating CRH is mediated by binding to CRH receptor, *g*_c_(*c*) will be chosen to be a Hill-type function, with Hill-exponent *n*, similar in form to the function *gR*(*OR*) used in Eqs. 13 and 18. The parameter 1 – *μ*_c_ represents the basal release rate of CRH relative to the maximum release rate and 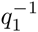 represents the normalized CRH level at which the positive effect is at half-maximum.

### Fast-slow variable separation and bistability

Since we assume the negative feedback effect of cortisol on synthesis of CRH operates over the longest characteristic timescale *t*_c_ in the problem, the full model must be studied across two separate timescales, a *fast timescale t*, and a *slow timescale τ* = *t*/*t*_c_ ≡ *εt*. The full model (Eqs. 14-18) can be succinctly written in the form

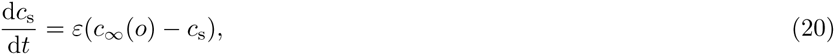

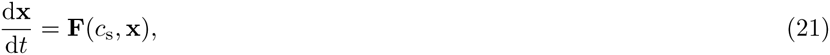

where **x** = (*c*, *a*, *o*, *r*) is the vector of fast dynamical variables, and **F**(*c*_s_, **x**) denotes the right-hand-sides of Eqs. 15-18. We refer to the fast dynamics described by d**x**/d*t* = **F**(*c*_s_, **x**) as a *fast flow*. In the *ε* → 0 limit, it is also easy to see that to lowest order *c*_s_ is a constant across the fast timescale and is a function of only the slow variable *τ*.

Under this timescale separation, the first component of Eq. 21(Eq. 15) can be written as

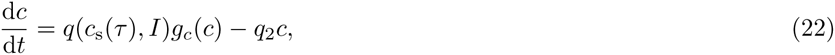

where *q*(*c*_s_(*τ*),*I*) ≡ *q*_0_*Ih*(*c*_s_(*τ*)) = *q*_0_*I*(1 – *e*^‒*kc*_s_(*τ*^)) is a function of *c*_s_(*τ*) and *I*. Since *c*_s_ is a function only of the slow timescale *τ*, *q* can be viewed as a bifurcation parameter controlling, over short timescales, the fast flow described by Eq. 22. Once *c*(*t*) quickly reaches its non-oscillating quasi-equilibrium value defined by d*c*/d*t* = *qg*_*c*_(*c*) – *q2*^*c*^ = 0, it can be viewed as a parametric term in Eq. 16 of the pituitary-adrenal (PA) subsystem.

Due to the nonlinearity of *g*_c_(*c*), the equilibrium value *c*(*q*) satisfying *qg*_*c*_(*c*) = *q*_2_^*c*^ may be multi-valued depending on *q*, as shown in Figs. 2A and 2B. For certain values of the free parameters, such as *n*, 1 – *μ*_c_, and *q*_1_, bistability can emerge through a saddle-node bifurcation with respect to the bifurcation parameter *q*. Fig. 2.B shows the bifurcation diagram, *i.e*., the nullcline of *c* defined by *qg*_*c*_(*c*) = *q*_2_^*c*^.

**Figure 2:**
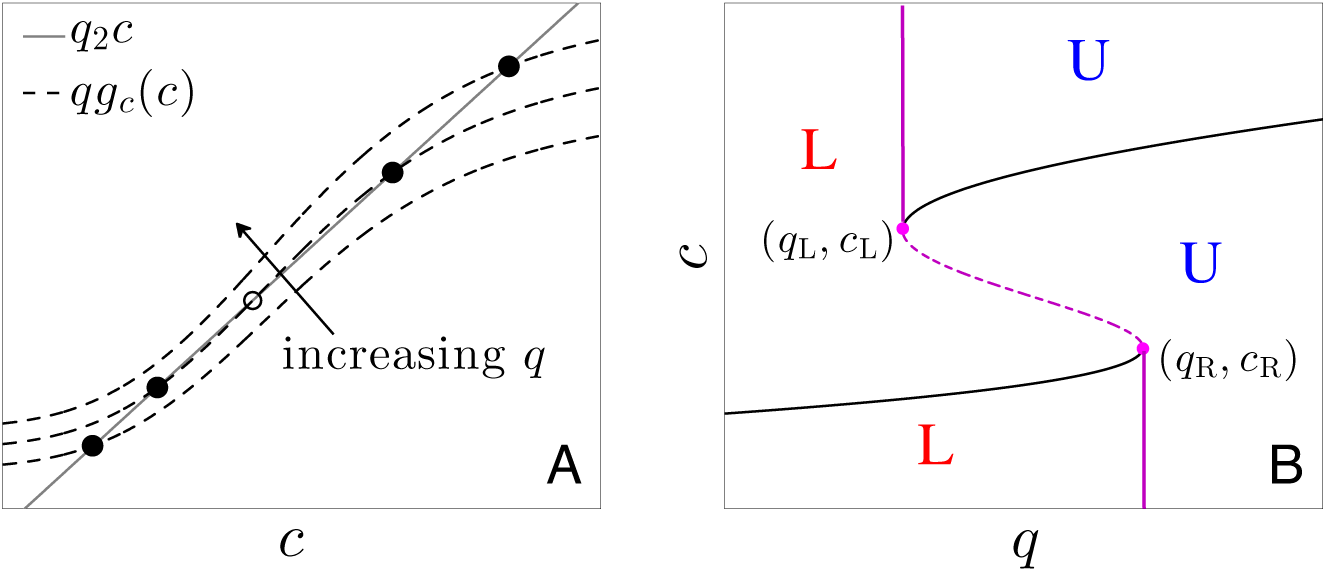
Nonlinear *g*_*c*_(*C*) and bistability of fast variables. (A) The stable states of the decoupled system in Eq. 22 can be visualized as the intersection of the two functions *qg*_*c*_(*c*) (dashed curve) and *q*2^*C*^ (gray line). For a given Hill-type function *g*_c_(*c*), Eq. 22 can admit one or two stable states (solid circles), depending on function parameters. The unstable steady state is indicated by the open circle. (B) Bifurcation diagram of the decoupled system (Eq. 22) with *q* as the bifurcation parameter. Solid and dashed segments represent stable and unstable steady states of the fast variables, respectively. *L* and *U* label basins of attraction associated with the lower and upper stable branches of the *c*-nullcline. Left and right bifurcation points (*q*_*L*_,*c*_*L*_) and (*q*_*R*_,*c*_*R*_) are indicated. Fixed points of c appear and disappear through saddle node bifurcations as *q* is varied through *q*_*L*_ and *q*_*R*_.

For equilibrium values of *c* lying within a certain range, the PA-subsystem can exhibit a limit cycle in (*a*,*o*,*r*) [13] that we express as (*a*^*^(*θ*; *c*),*o*^*^(*θ*; *c*;),*r*^*^(*θ*; *c*)), where *θ* = 2π*t*/*t*_*p*_ (*c*) is the phase along the limit cycle. The dynamics of the PA-subsystem depicted in Fig. 3. indicate the range of *c* values that admit limit cycle behavior for (*a*, *o*, *r*), while the fast *c*-nullcline depicted in Fig. 2.B restricts the range of bistable *c* values. Thus, bistable states that also support oscillating (*a*, *o*, *r*) are possible only for values of *c* that satisfy both criteria.

**Figure 3:**
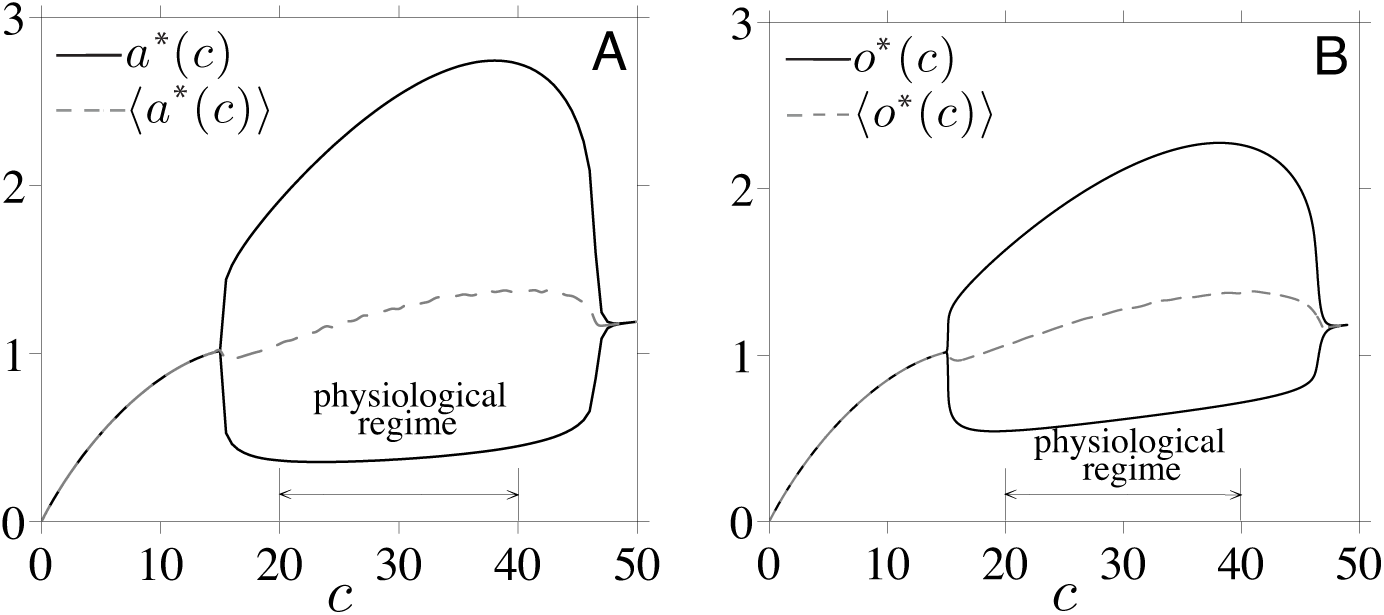
Dynamics of the oscillating PA-subsystem as a function of fixed *c*. (A)Maximum/minimum and period-averaged values of ACTH, *a*(*T*), as a function of circulating CRH. (B) Maximum/minimum and period-averaged values of cortisol *o*(*t*). Within physiological CRH levels, ACTH, GR (not shown), and cortisol oscillate. The minimum, maximum, and period-averaged cortisol levels typically increase with increasing *c*. The plot was generated using dimensionless variables *c*, *a*, and *o* with parameter values specified in [43] and *t*_d_ = 1.44, corresponding to a delay of *T*_d_ = 15min.

Since in the *ε* → 0 limit, circulating CRH only feeds forward into *a*, *o*, and *r*, a complete description of all the fast variables can be constructed from just *c* which obeys Eq. 22. Therefore, to visualize and approximate the dynamics of the full five-dimensional model, we only need to consider the 2D projection onto the fast *c* and slow *c*_s_ variable. A summary of the time-separated dynamics of the variables in our model is given in Fig. 4..

**Figure 4:**
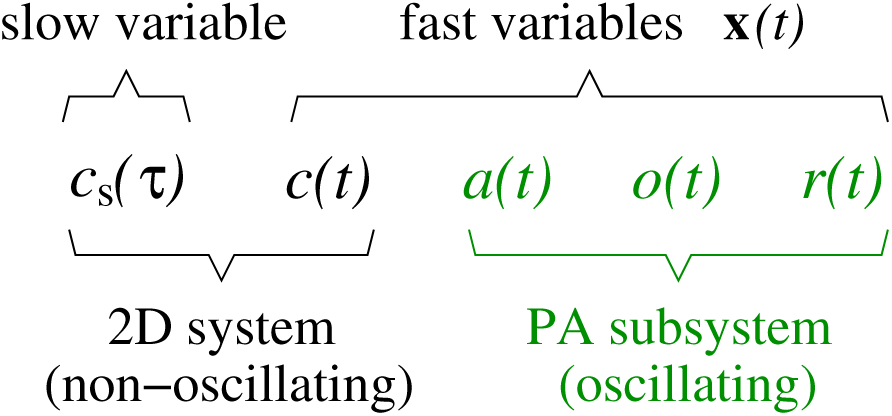
Classification of variables. Variables of the full five-dimensional model are grouped according to their dynamical behavior. *C*_s_(*τ*) is a slow variable, while **x**(*t*) = (*c*, *a*, *o*, *r*) are fast variables. Of these, (*a*, *o*, *r*) form the typically oscillatory PA-subsystem that is recapitulated by c. In the *ε* = 1/*t*_*c*_ ≪ 1 limit, the variable *C*_s_(*τ*) slowly relaxes towards a period-averaged value 〈*c*_∞_(*o*(*c*))〉. Therefore, the full model can be accurately described by its projection onto the 2D (*c*_s_, *c*) phase space.

To analyze the evolution of the slow variable *c*_s_(*τ*), we write our equations in terms of *τ* = *εt*:

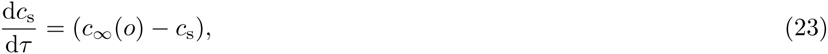

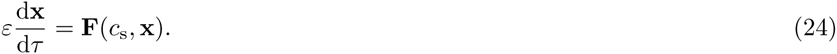

In the *ε* → 0 limit, the “outer solution” **F**(*c*_s_, **x**) ≈ 0 simply constrains the system to be on the fast *c*-nullcline defined by *qg*_*c*_(*c*) = *q*_2_^*c*^. The slow evolution of *c*_s_(*τ*) along the fast *c*-nullcline depends on the value of the fast variable *o*(*t*) through *c*_∞_ (*o*). To close the slow flow subsystem for *c*_s_(*τ*), we fix *c* to its equilibrium value as defined by the fast subsystem and approximate *c*_∞_(*o*(*c*)) in Eq. 23 by its period-averaged value

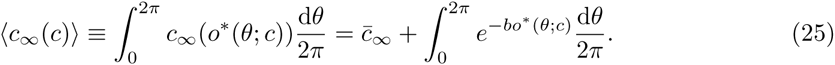

Since *o*^*^ increases with *c*, 〈*c*_∞_(*c*)〉 is a decreasing function of *c* under physiological parameter regimes. This period-averaging approximation allows us to relate the evolution of *c*_s_(*τ*) in the slow subsystem directly to *c*. The evolution of the slow subsystem is approximated by the closed (*c*_s_, *c*) system of equations

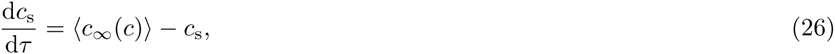

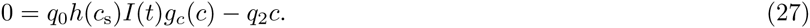

with 〈*c*_∞_(*c*)〈 evaluated in Eq. 25. By self-consistently solving Eqs. 26 and 27, we can estimate trajectories of the full model when they are near the *c*-nullcline in the 2D (*c*_s_, *c*)-subsystem. We will verify this in the following section.

### Nullcline structure and projected dynamics

The separation of timescales results in a natural description of the fast *c*-nullcline in terms of the parameter *q* (Fig. 2.) and the slow *c*_s_-nullcline (defined by the relation *c*_s_ = 〈*c*_∞_(*c*)〈 relating *c*_s_ to *c*) in terms of *c*. However, the *c*-nullcline is plotted in the (*q*, *c*)-plane while the *c*_s_-nullcline is defined in the (*c*, *c*_s_)-plane. To plot the nullclines together, we relate the equilibrium value of *c*_s_, 〈*c*_∞_(*c*)〉, to the *q* coordinate through the monotonic relationship *q*(*c*_s_) = *q*_0_*Ih*(〈*c*_∞_(*c*)〉) = *q*_0_*I*(1 – *e*^‒*k*〈*c*_∞_(*c*)〉^) and transform the *c*_s_ variable into the *q* parameter so that both nullclines can be plotted together in the (*q*, *c*)-plane. These transformed *c*s-nullclines will be denoted “*q*-nullclines.”

We assume a fixed basal stress input *I* = 1 and plot the *q*-nullclines in Fig. 5.A for increasing values of *k*, the parameter governing the sensitivity of CRH release to stored CRH. From the form *h*(〈*c*_∞_(*c*)〉) = (1 – *e*^*–k*〈*c*_∞_(*c*)〉^), both the position and the steepness of the *q*-nullcline in (*q*, *c*)-space depend strongly on *k*. Fig. 5.B shows a fast *c*-nullcline and a slow *q*-nullcline (transformed *c*s-nullcline) intersecting at both stable branches of the fast *c*_s_-nullcline. Here, the flow field indicates that the 2D projected trajectory is governed by fast flow over most of the (*q*, *c*)-space.

**Figure 5:**
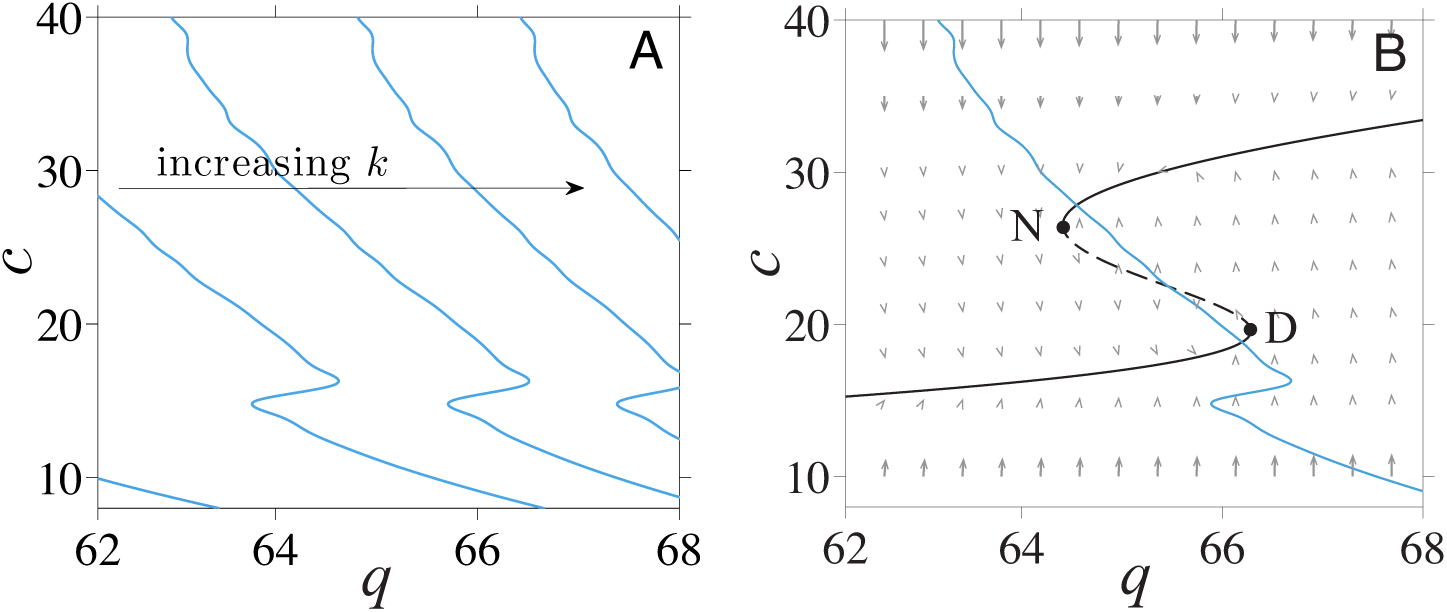
Slow and fast nullclines and overall flow field. (A) The nullcline of *C*_s_ in the *ε* → 0 limit is defined by *C*_s_ = 〈*c*_∞_(*c*)〉. To plot these slow nullclines together with the fast *c*-nullclines, we transform the variable *C*_s_ and represent it by *q* through the relation *q* = *q0*^*h*^(*C*_s_). These transformed nullclines then become a function of c and can be plotted together with the fast *c*-nullclines. For each fixed value of *c*, *o*(*t*; *c*) is computed by employing a built-in DDE solver dde23 in MATLAB. The numerical solution is then used to approximate 〈*c*_∞_(*c*)〉 in Eq. 25 by Euler’s method. The q-nullcline shifts to the right and gets steeper as *k* increases. (B) The fast *c*-nullcline defined by *qg*_C_(*C*) = *q2*^*C*^ (black curve) is plotted together with the slow *C*_s_-nullcline plotted in the (*q*, *c*) plane (“*q*-nullcline,” blue curve). Here, two intersections arise corresponding to a high-cortisol normal (N) stable state and a low-cortisol diseased (D) stable state. The flow vector field is predominantly aligned with the fast directions toward the *c*-nullcline.

How the fast and slow nullclines cross controls the long-term behavior of our model in the small *ε* limit. In general, the number of allowable nullcline intersections will depend on input level *I* and on parameters (*q*_0_,…,*p*_6_, *b*, *k*, *n*, *μ*_*c*_, *t*_d_). Other parameters such as *q*_0_, *q*_1_, and *μ*_*c*_ appear directly in the fast equation for *c* and thus most strongly control the fast *c*-nullcline. Fig. 6.A shows that for a basal stress input of *I* = 1 and an intermediate value of *k*, the nullclines cross at both stable branches of the fast subsystem. As expected, numerical simulations of our full model show the fast variables (*a*, *o*, *r*) quickly reaching their oscillating states defined by the *c*-nullcline while the slow variable *q* = *q*_0_*Ih*(*c*_s_) remains fairly constant. Independent of initial configurations that are not near the *c*-nullcline in (*q*, *c*)-space, trajectories quickly jump to one of the stable branches of the *c*-nullcline with little motion towards the *q*-nullcline, as indicated by *ξ*f in Fig. 6.A.

**Figure 6:**
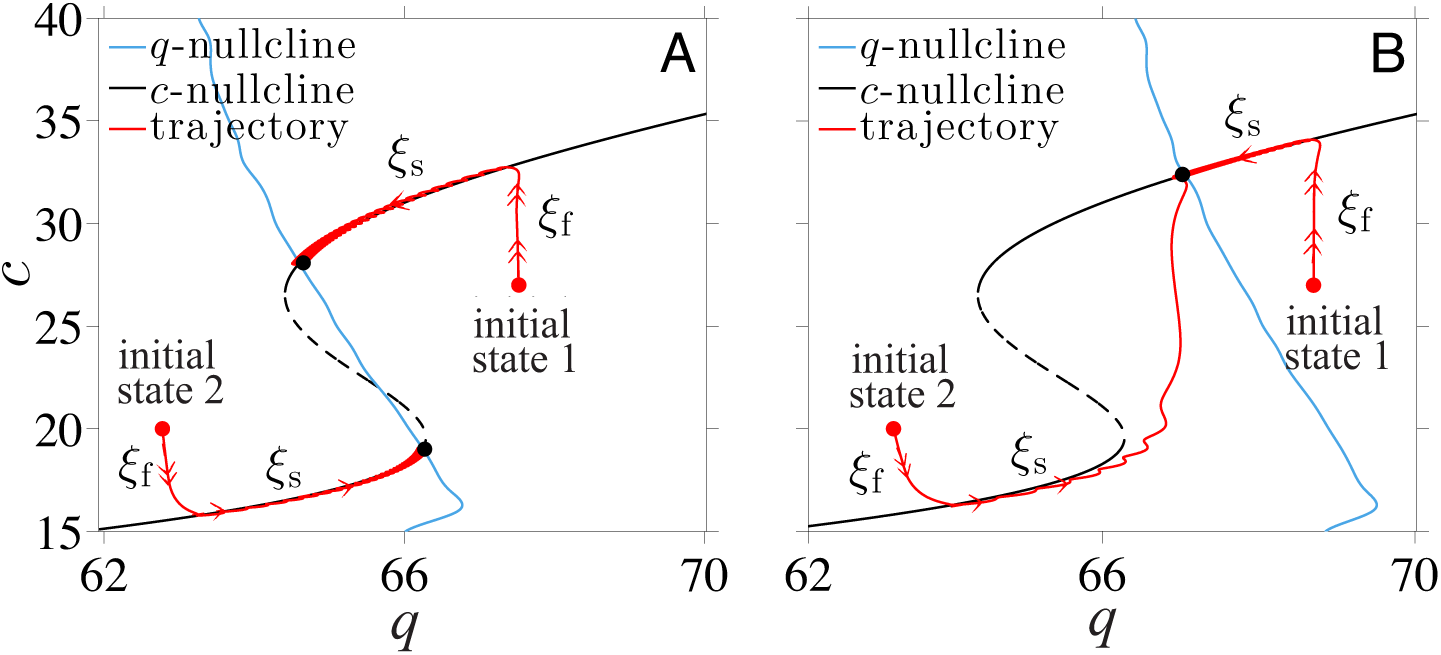
Equilibria at the intersections of nullclines. (A) For intermediate values of *k*, there are three intersections, two of them representing stable equilibria. Solid red lines are projections of two trajectories of the full model, with initial states indicated by red dots and final stable states shown by black dots. The full trajectories approach the intersections of the *q*-nullcline (blue) and *c*-nullcline (black). (B) For large *k* there is only one intersection at the upper branch of the *c*-nullcline. Two trajectories with initial states near different branches of the *c*-nullcline both approach the unique intersection (black dot) on the upper branch. The scenario shown here corresponds to a Type *I* nullcline structure as described in the Additional File.

Once near the *c*-nullcline, say when |**F**(*c*_s_, **x**)| ≪ ε, the trajectories vary slowly according to Eqs. 23. Here, the slow variable *c*_s_ relaxes to its steady state value while satisfying the constraint **F**(*c*_s_, **x**) ≈ 0. In (*q*,*c*)–space, the system slowly slides along the *c*-nullcline towards the *q*-nullcline (the *ξ*_s_ paths in Fig. 6.A). This latter phase of the evolution continues until the system reaches an intersection of the two nullclines, indicated by the filled dot, at which the reduced subsystem in *c*_s_ and *c* reaches equilibrium.

For certain values of *k* and if the fast variable *c* is bistable, the two nullclines may intersect within each of the two stable branches of the *c*-nullcline and yield the two distinct stable solutions shown in Fig. 6.A. For large *k*, the two nullclines may only intersect on one stable branch of the *c*-nullcline as shown in Fig. 6.B. Trajectories that start within the basin of attraction of the lower stable branch of the *c*-nullcline (“initial state 2” in Fig. 6.B) will stay on this branch for a long time before eventually sliding off near the bifurcation point and jumping to the upper stable branch. Thus, the long-term behavior of the full model can be described in terms of the locations of the intersections of nullclines of the reduced system.

## Results and Discussion

The dual-nullcline structure and existence of multiple states discussed above results from the separation of slow CRH synthesis process and fast CRH secretion process. This natural physiological separation of time scales ultimately gives rise to slow dynamics along the fast *c*-nullcline during stress. The extent of this slow dynamics will ultimately determine whether a transition between stable states can be induced by stress. In this section, we explore how external stress-driven transitions mediated by the fast-slow negative feedback depend on system parameters.

Changes in parameters that accompany trauma can lead to shifts in the position of the nullclines. For example, if the stored CRH release process is sufficiently compromised by trauma (smaller *k*), the slow *q*-nullcline moves to the left, driving a bistable or fully resistant organism into a stable diseased state. Interventions that increase *k* would need to overcome hysteresis in order to restore normal HPA function. More permanent changes in parameters are likely to be caused by physical rather than by psychological traumas since such changes would imply altered physiology and biochemistry of the person. Traumatic brain injury (TBI) is an example of where parameters can be changed permanently by physical trauma. The injury may decrease the sensitivity of the pituitary to cortisol-GR complex, which can be described by decreasing *p*_2_ in our model. Such change in parameter would lead to a leftward shift of the *q*-nullcline and an increased likelihood of hypocortisolism.

In the remainders of this work, we focus on how external stress inputs can by themselves induce stable but reversible transitions in HPA dynamics *without* changes in physiological parameters. Specifically, we consider only temporary changes in *I*(*t*) and consider the time-autonomous problem. Since the majority of neural circuits that project to the PVN are excitatory [44], we assume external stress stimulates CRH neurons to release CRH above its unit basal rate and that *I*(*t*) = 1 + *I*_ext_(*t*)(I_base_ = 1) with *I*_ext_ ≥ 0.

To be more concrete in our analysis, we now choose our nullclines by specifying parameter values. We estimate the values of many of the dimensionless parameters by using values from previous studies, as listed in Table S1 in the Additional File. Of the four remaining parameters, *μ*_*c*_, *q*_0_, *q*_1_, and *k*, we will study how our model depends on *k* while fixing *μ*_*c*_*q*_0_, and *q*_1_. Three possible nullcline configurations arise according to the values of *μ*_*c*_, *q*_0_, and *q*_1_ and are delineated in the Additional File. We have also implicitly considered only parameter regimes that yield oscillations in the PA subsystem at the stable states defined by the nullcline intersections.

Given these considerations, we henceforth chose *μ*_*c*_ = 0.6, *q*_1_ = 0.04, and *q*_0_ = 77.8 for the rest of our analysis. This choice of parameters is motivated in the Additional File and corresponds to a so-called “Type I” nullcline structure. In this case, three possibilities arise: one intersection on the lower stable branch of the *c*-nullcline if *k* < *k*_L_, two intersections if *k*_L_ < *k* < *k*_R_ (Fig. 6.A), and one intersection on the upper stable branch of the *c*-nullcline if *k* > *k*_R_ (Fig. 6.B). For our chosen set of parameters and a basal stress input *I* = 1, the critical values *k*_L_ = 2.5 < *k*_R_ = 2.54 are given by Eq. A3 in the Additional File.

## Normal stress response

Activation of the HPA axis by acute stress culminates in an increased secretion of all three main hormones of the HPA axis. Persistent hypersecretion may lead to numerous metaboli*c*, affective, and psychotic dysfunctions [45, 46]. Therefore, recovery after stress-induced perturbation is essential to normal HPA function. We explore the stability of the HPA axis by initiating the system in the upper of the two stable points shown in Fig. 7.A and then imposing a 120min external stress input *I*_*ext*_ = 0.1. The HPA axis responds with an increase in the peak level of cortisol before relaxing back to its original state after the stress is terminated (Fig. 7.B). This transient process is depicted in the projected (*q*, *c*)-space in Fig. 7.A.

**Figure 7:**
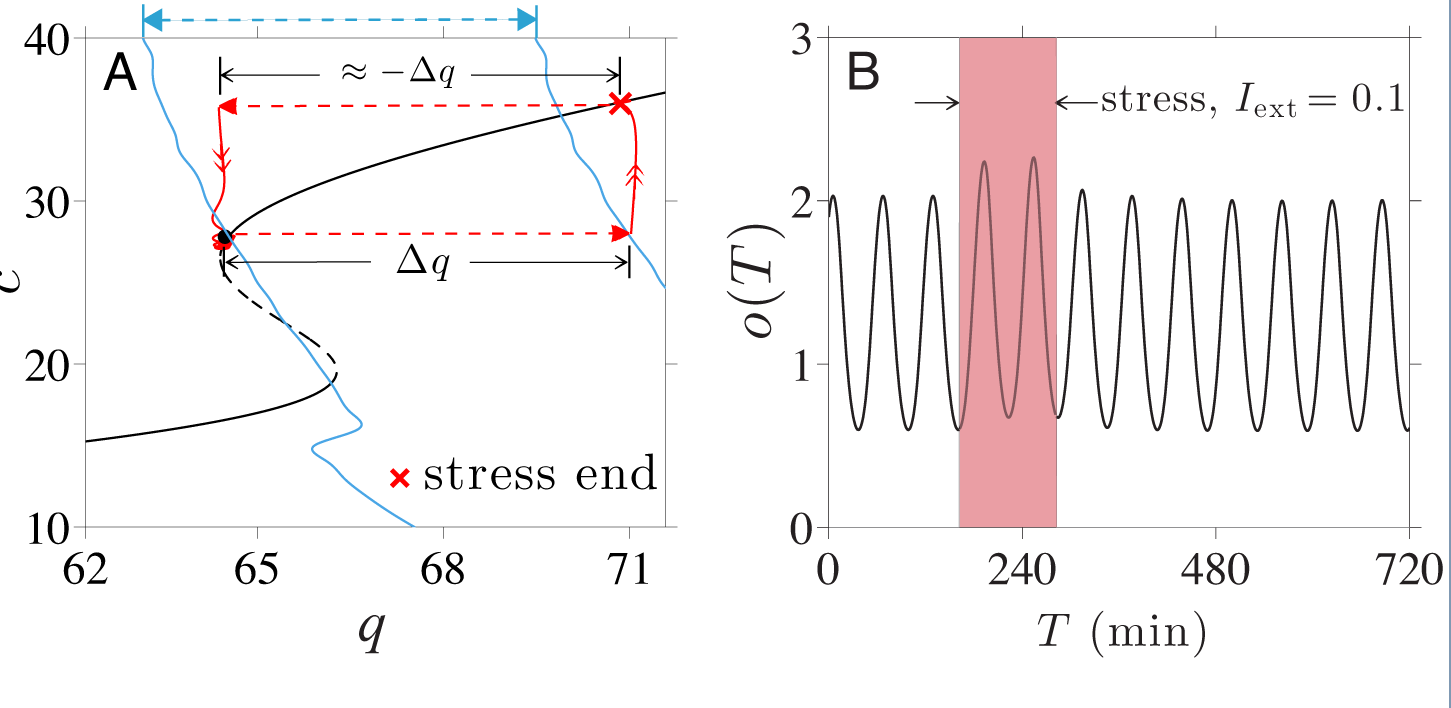
Normal stress response. Numerical solution for the response to a 120min external stress *I*_ext_ = 0.1. (A) At the moment the external stress is turned on, the value of (*q*,*C*) increases from its initial stable solution at (64.4, 27) to (71, 27) after which the circulating CRH level *C*, quickly reaches the fast *c*-nullcline (black) before slowly evolving along it towards the slow *q*-nullcline (blue). After short durations of stress, the system returns to its starting point within the normal state basin. (B) The peaks of the cortisol level are increased during stress (red) but return to their original oscillating values after the stress is turned off.

Upon turning on stress, the lumped parameter *q* and the slow nullcline shift to the right by 10% since *q* = qo(1+*I*_ext_)*h*(〈*c*_∞_(*c*)〉) (see Fig. 7.A). The trajectory will then move rapidly upward towards the new value of *c* on the *c*-nullcline; afterwards, it moves very slowly along the *c*-nullcline towards the shifted *q*-nullcline. After 120min, the system arrives at the “x” on the *c*-nullcline (Fig. 7.A). Once the stress is shut off the *q*-nullcline returns to its original position defined by *I* = 1. The trajectory also jumps back horizontally to near the initial *q* value and subsequently quickly returns to the original upper-branch stable point.

### External stress induces transition from normal to diseased state

We now discuss how transitions from a normal to a diseased state can be induced by *positive* (excitatory) external stress of sufficient duration. In Fig. 8., we start the system in the normal high*-c* state.

**Figure 8:**
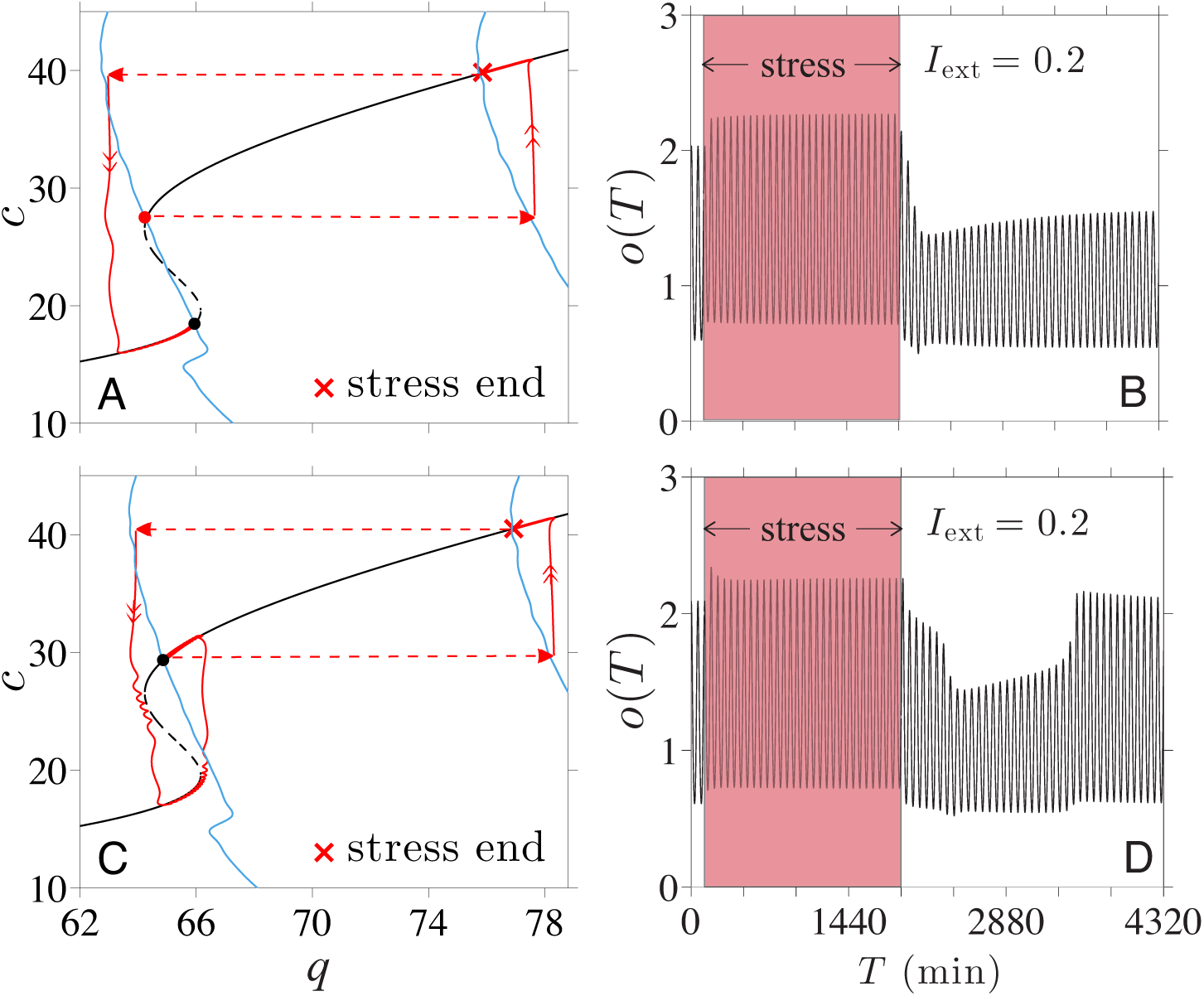
Stress-induced transitions into an oscillating low-cortisol diseased state. An excitatory external stress *I*_ext_ = 0.2 is applied for 30hrs. Here, the system reaches the new stable point set by *I* = 1.2 before stress is terminated and the *q*-nullcline reverts to its original position set by *I* = 1. (A) At intermediate values of 2.5 < *k* < 2.54, when two stable state arise, a transition from the normal high-cortisol state into the diseased low-cortisol state can be induced by chronic external stress. (B) Numerical solutions of cortisol level *o*(*T*) plotted against the original time variable *T* shows the transition to the low-cortisol diseased state shortly after cessation of stress. (*c*) and (D) If *k* > *k*_R_ = 2.54, only the normal stable state exists. The system will recover and return to its original healthy state after a transient period of low cortisol.

Upon stimulation of the CRH neurons through *I*_ext_ > 0, both CRH and average glucocorticoid levels are increased while the average value of *c*_∞_(*o*(*t*)) is decreased since *c*_∞_(*o*) is a decreasing function of *o*. As *c*_s_(*τ*) slowly decays towards the decreased target value of 〈*c*_∞_(*o*(*c*))〉, *h*(*c*_s_(*τ*)), and hence *q*(*c*_s_), also decrease. As shown in Fig. 8.A, much of this decrease occurs along the high-*c* stable branch of the *c*-nullcline. Once the external stress is switched off, *q* will jump back down by a factor of 1/(1 + *I*_ext_). If the net decrease in *q* is sufficient to bring it below the bifurcation value *q*_*L*_ ≈ 64 at the leftmost point of the upper knee, the system crosses the separatrix and approaches the alternate, diseased state. Thus, the normal-to-diseased transition is more likely to occur if the external stress is maintained long enough to cause a large net decrease in *q*, which includes the decrease in *q* incurred during the slow relaxation phase, plus the drop in *q* associated with cessation of stress. The minimum duration required for normal-to-diseased transition should also depend on the magnitude of *I*_ext_. The relation between the stressor magnitude and duration will be illustrated in the Additional Files.

A numerical solution of our model with a 30hr *I*_ext_ = 0.2 was performed, and the trajectory in (*q*, *c*)-space is shown in Fig. 8.A. The corresponding cortisol level along this trajectory is plotted in Fig. 8.B, showing that indeed a stable transition to the lower cortisol state occurred shortly after the cessation of stress. In addition to a long-term external stress, the stable transition to a diseased state requires 2.5 < *k* < 2.54 and the existence of two stable points. On the other hand, when *k* > *k*_R_ = 2.54, the enhanced CRH release stimulates enough cortisol production to drive the sole long term solution to the stable upper normal branch of the *c*-nullcline, rendering the HPA system *resistant* to stress-induced transitions.

The response to chronic stress initially follows the same pattern as described above for the two-stable-state case, as shown in Fig. 8.C. However, the system will continue to evolve along the lower branch towards the *q*-nullcline, eventually sliding off the lower branch near the right bifurcation point (indicated in Fig. S2 by (*q*_*R*_, *c*_*R*_)) and returning to the single normal equilibrium state. Thus, when *k* is sufficiently high, the system may experience a transient period of lowered cortisol level after chronic stress but will eventually recover and return to the normal cortisol state. The corresponding cortisol level shown in Fig. 8.D shows this recovery at *T* ≈ 3400min, which occurs approximately 1500min after the cessation of stress.

### Transition to diseased state depends on stress timing

We have shown how transitions between the oscillating normal and diseased states depend on the duration of the external stress *I*_ext_. However, whether a transition occurs also depends on the *time* – relative to the phase of the intrinsic ultradian oscillations – at which a fixed-duration external stress is initiated. To illustrate this dependence on phase, we plot inFigs. 9A and B two solutions for *o*(*t*) obtained with a 250min *I*_ext_ = 0.1 initiated at different phases of the underlying cortisol oscillation. If stress is initiated during the rising phase of the oscillations, a transition to the low-cortisol diseased state occurs and is completed at approximately *t* = 1000min (Fig. 9.A,C). If, however, stress is initiated during the falling phase, the transition does not occur and the system returns to the normal stable state (Fig. 9.B, D). In this case, a longer stress duration would be required to push the trajectory past the low-q separatrix into the diseased state.

**Figure 9:**
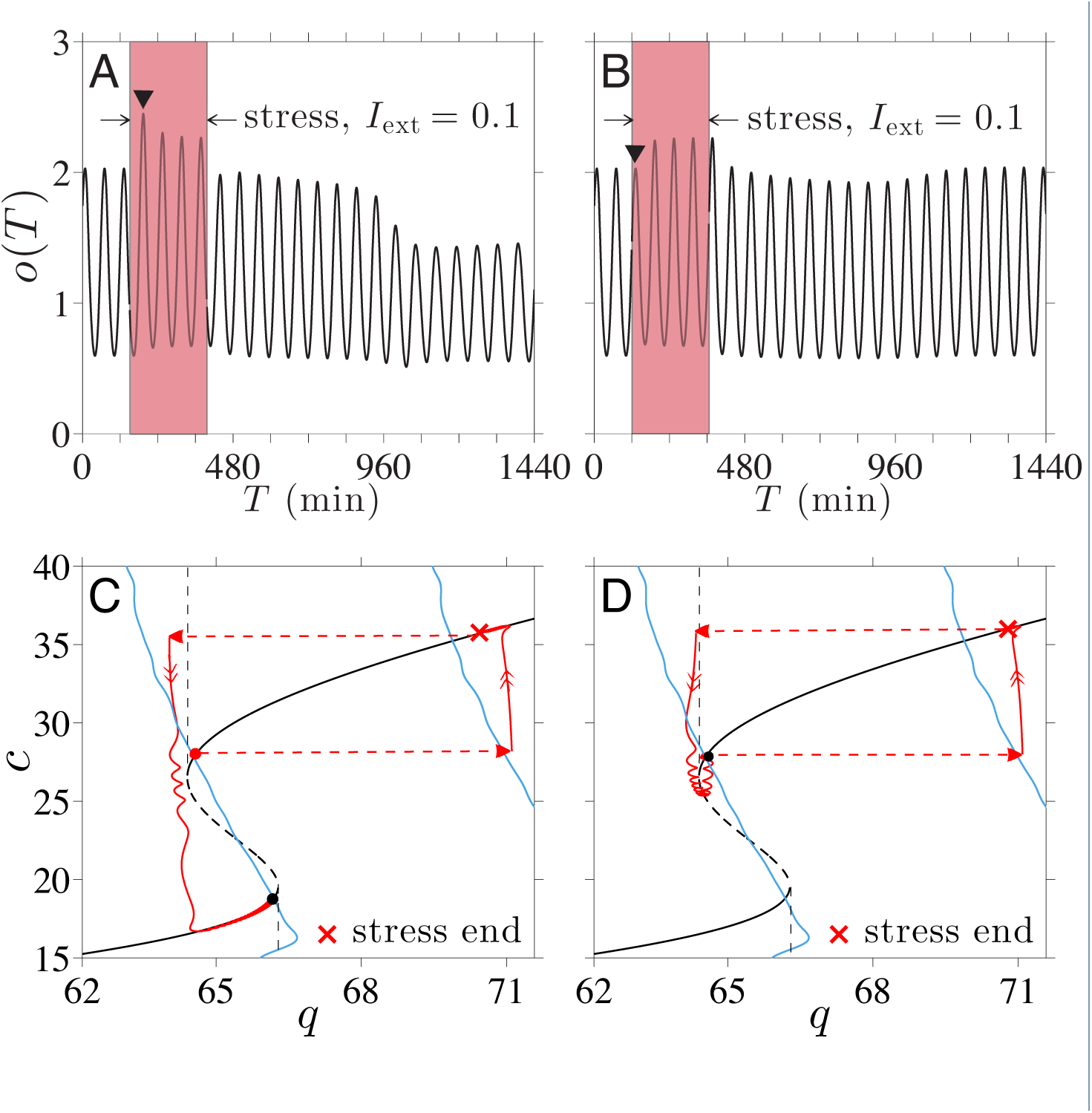
Stress timing and transition to low-cortisol oscillating state. Cortisol levels in response to *I*_ext_ = 0.1 applied over 250min. (A) If stress is initiated at *T* = 150min, a transition to the low-cortisol diseased state is triggered. (B) If stress is initiated at *T* = 120min, the system returns to its normal high-cortisol state. Note that the first peak (marked by “▼”) during the stress in (A) is higher than the first peak in (B). (*c*) If stress is initiated at *T* = 150min, stress cessation and the slow relaxation along the *c*-nullcline during stress are sufficient to bring *q* just left of the separatrix, inducing the transition. (D) For initiation time *T* = 120min, *q* remains to the right of the separatrix, precluding the transition.

As discussed earlier, an increase in period-averaged cortisol level during stress drives a normal-to-diseased state transition. We see that the period-averaged level of cortisol under increased stress is different for stress started at 120min from stress started at 150min. As detailed in the Additional File, the amplitude of the first cortisol peak after the start of stress is significantly lower when the applied stress is started during the falling phase of the intrinsic cortisol oscillations. The difference between initial responses in *o*(*t*) affects the period-averaging in 〈*c*_∞_(*o*)〉 during external stress, ultimately influencing *c*_s_ and consequently determining whether or not a transition occurs. Note that this phase dependence is appreciable only when stress duration is near the threshold value that brings the system close to the separatrix between normal and diseased basins of attraction. Trajectories that pass near separatrices are sensitive to small changes in the overall negative feedback of cortisol on CRH synthesis, which depend on the start time of the stress signal.

### Stress of intermediate duration can induce “reverse” transitions

We can now use our theory to study how *positive* stressors *I*_ext_ may be used to induce “reverse” transitions from the diseased to the normal state. Understanding these reverse transitions may be very useful in the context of exposure therapy (ET), where PTSD patients are subjected to stressors in a controlled and safe manner, using for example, computer-simulated “virtual reality exposure.” Within our model we can describe ET as external stress ( *I*_ext_ > 0) applied to a system in the stable low-*c* diseased state. The resulting horizontal shift in *q* causes the system to move rightward across the separatrix and suggests a transition to the high-*c* normal state can occur upon termination of stress. As shown in Fig. 10.A, if stressor of sufficient duration is applied, the trajectory reaches a point above the unstable branch of the *c*-nullcline upon termination leading to the normal, high-cortisol state (Fig. 10.B). Since the initial motion is governed by fast flow, the minimum stress duration needed to incite the diseased-to-normal transition is short, on the timescale of minutes. However, if the stressor is applied for too long, a large reduction in *q* is experienced along the upper stable branch. Cessation of stress might then lower *q* back into the basin of attraction of the low-cortisol diseased state (Fig. 10.C). Fig. 10.D shows the cortisol level transiently increasing to a normal level before reverting back to low levels after approximately 1400min.

**Figure 10:**
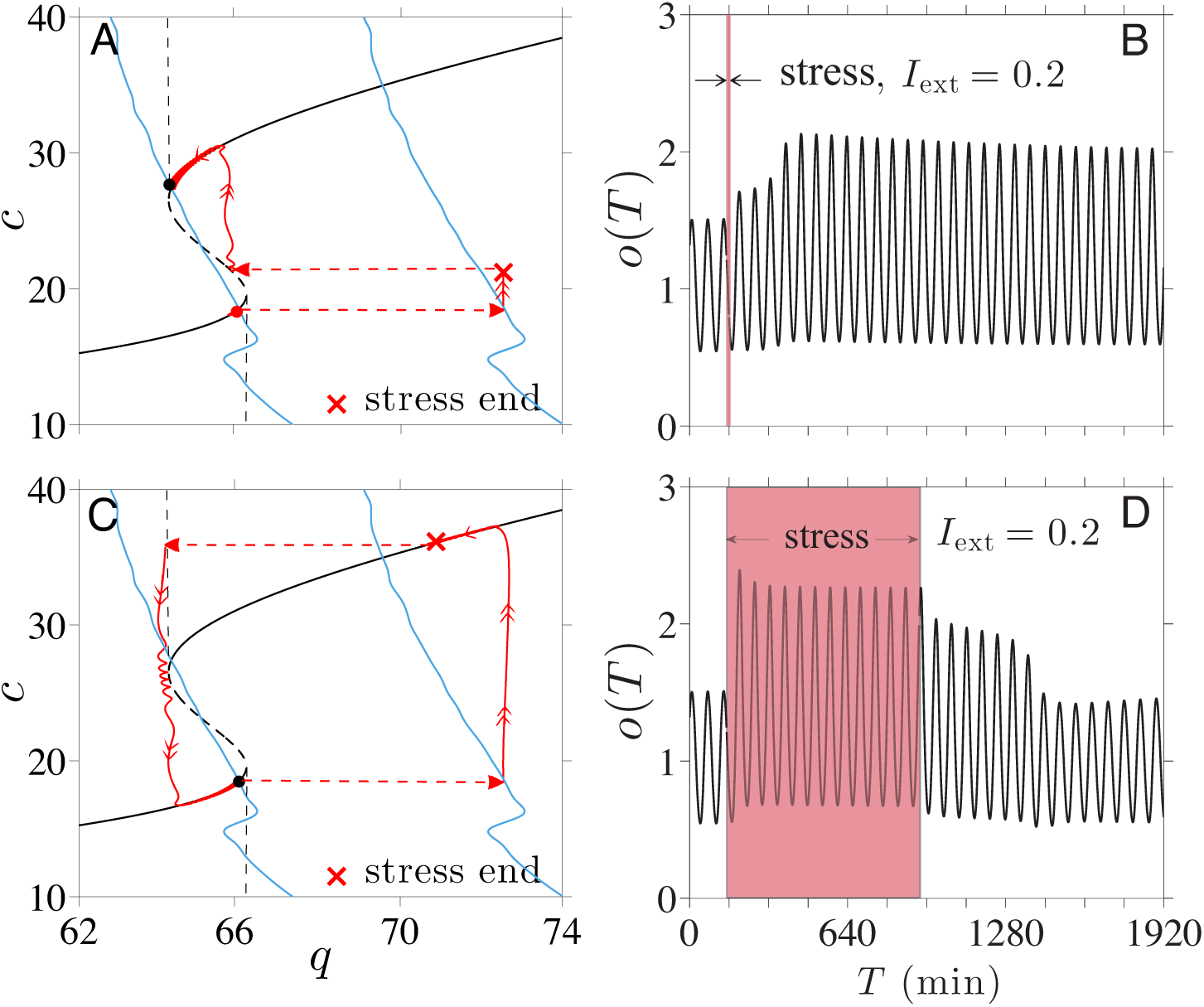
Stress-induced transitions to high-cortisol oscillating state. (A) Projected 2D system dynamics when a stressor of amplitude *I*_ext_ = 0.1 is applied for 9min starting at *T* = 120min. c is increased just above the unstable branch (*c* ≈ 20) to allow the unstressed system to cross the separatrix and transition to the normal high-*c* stable state. (B) The plot of *o*(*T*) shows the transition to the high-cortisol, high-oscillation amplitude state shortly after the 9min stress. (c) A stressor turned off after 780min (13hrs) leaves the system in the basin of attraction of the diseased state. (D) Cortisol levels are pushed up but after about 1400min relax back to levels of the original diseased state.

Within our dynamical model, stresses need to be of *intermediate* duration in order to induce a stable transition from the diseased to the normal state. The occurrence of a reverse transition may also depend on the phase (relative to the intrinsic oscillations of the fast PA subsystem) over which stress was applied, especially when the stress duration is near its transition thresholds. For a reverse diseased-to-normal transition to occur, the decrease in *c*s cannot be so large that it brings the trajectory past the left separatrix, as shown in Fig. 10.C. Therefore, near the maximum duration, stress initiated over the falling phase of cortisol oscillation will be more effective at triggering the transition to a normal high-cortisol state. Overall, these results imply that exposure therapy may be tuned to drive the dynamics of the HPA axis to a normal state in patients with hypocortisolism-associated stress disorders.

## Summary and Conclusions

We developed a theory of HPA dynamics that includes stored CRH, circulating CRH, ACTH, cortisol and glucocorticoid receptor. Our model incorporates a fast self-upregulation of CRH release, a slow negative feedback effect of cortisol on CRH synthesis, and a delay in ACTH-activated cortisol synthesis. These ingredients allow our model to be separated into slow and fast components and projected on a 2D subspace for analysis.

Depending on physiological parameter values, there may exist zero, one, or two stable simultaneous solutions of both fast and slow variables. For small *k*, CRH release is weak and only the low-CRH equilibrium point arises; an individual with such *k* is trapped in the low-cortisol “diseased” state. For large *k*, only the high-CRH normal state arises, rendering the individual resistant to acquiring the long-term, low-cortisol side-effect of certain stress disorders. When only one stable solution arises, HPA dysregulation must depend on changes in parameters resulting from permanent physiological modifications due to *e.g*., aging, physical traum*a*, or stress itself [46, 47]. For example, it has been observed that older rats exhibit increased CRH secretion while maintaining normal levels of CRH mRNA in the PVN [48]. Such a change could be interpreted as an age-dependent increase in *k*, which, in our model, implies that aging makes the organism more resistant to stress-induced hypocortisolism. Indeed, it has been suggested that prevalence of PTSD declines with age [49, 50].

Other regulatory systems that interacts with or regulate the HPA axis can also affect parameter values in our model. Gonadal steroids, which are regulated by another neuroendocrine system called the hypothalamic-pituitary-gonadal (HPG) axis, activate the preoptic area (POA) of the hypothalamus [51, 52], which in turn attenuates the excitatory effects of medial amygdala stimulation of the HPA axis [53]. Thus, low testosterone levels associated hypogonadism would effectively increase *I*( *t*) within our model, shift the *q*-nullcline in the ( *q*, *c*)-space, and in turn increase cortisol levels. One might consider this as a possible explanation for chronically elevated cortisol levels observed in major depressive disorder patients who suffers from hypogonadism. Although it is beyond the scope of this paper, one may further investigate role of gonadal hormones, or role of any other interacting systems, in mediating stress response by considering which parameters would be affected in our model.

Within certain parameter regimes and for intermediate *k*, our theory can also exhibit bistability. When two stable solutions arise, we identify the states with low oscillating levels of cortisol as the diseased state associated with hypocortisolism. Transitions between different stable states can be induced by temporary external stress inputs, implying that HPA dysregulation may develop without permanent “structural” or physiological changes. Stresses that affect secretion of CRH by the PVN are shown to be capable of inducing transitions from normal to diseased states provided they are of sufficient duration (Fig. 8.).

Our model offers a mechanistic explanation to the seemingly counter-intuitive phenomenon of lower cortisol levels after *stress*-induced *activation* of cortisol production. Solutions to our model demonstrate that the negative-feedback effect of a temporary increase in cortisol on the synthesis process of CRH can slowly accumulate during the stress response and eventually shift the system into a different basin of attraction. Such a mechanism provides an alternative to the hypothesis that hypocortisolism in PTSD patients results from permanent changes in physiological parameters associated with negative-feedback of cortisol [54, 55].

We also find that external stress can induce the “reverse” transition from a diseased low-cortisol state to the normal high-cortisol state. Our results imply that re-exposure to stresses of *intermediate* duration can drive the system back to normal HPA function, possibly “decoupling” stress disorders from hypocortisolism.

Interestingly, we show that the minimum durations required for either transition depends on the time at which the stress is initiated relative to the phase of the intrinsic oscillations in (*a*, *o*, *r*). Due to subtle differences in cortisol levels immediately following stress initiation at different phases of the intrinsic cortisol oscillation, the different cumulative negative-feedback effect on CRH can determine whether or not a trajectory crosses a separatrix (Fig. 9.). When the duration of external stress is near its threshold, normal-to-diseased state transitions are easier to induce when stress is initiated during the rising phase of cortisol oscillations. Reverse diseased-to-normal transitions are more easily induced when stress is initiated during the falling phase.

In summary, our theory provides a mechanistic picture that connects cortisol dysregulation with stress disorders and a mathematical framework one can use to study the downstream effects of therapies such as brief eclectic psychotherapy (BEP) and exposure therapy (ET). Both therapies involve re-experiencing stressful situations directly or through imagination, and have been consistently proven effective as first-line treatments for PTSD symptoms [56–58]. Our results suggest that ET can directly alter and “decouple” the expression of cortisol from an underlying upstream disorder. Changes in neuronal wiring that typically occur over slower times scales is also expected after ET [59]. In our model, such changes would lead to slow changes in the basal input *I*(*t*). Thus, cortisol level may not be tightly correlated with PTSD, particularly in the context of ET.

It is important to emphasize that we modeled neuroendocrine dynamics downstream of the stress input *I*_ext_. How the form of the stress function *I*_ext_ depends on the type of stress experienced requires a more detailed study of more upstream processes, including how hormones might feedback to these higher-brain processes. Since *higher* cortisol levels are found among female PTSD patients with a history of childhood abuse [60] and among PTSD patients who have experienced a nuclear accident [61], future studies of such divergent, experience-dependent dysregulation will rely on more complex input functions *I*_ext_(*t*). For example, under periodic driving, complex resonant behavior should arise depending on the amplitude and frequency of the external stress *I*_ext_(*t*) and the nullcline structure of the specific system. Moreover, effects of other regulatory networks that interacts with the HPA axis can be included in our model through appropriate forms of *I*_ext_(*t*). For example, the effects of gonadal steroids in the stress response mentioned above can be further investigated by considering a form of *I*_ext_ (*t*) that is dependent on gonadal steroids level. Many other interesting properties, such as response to dexamethasone administration, can be readily investigated within our model under different system parameters.

## Author’s contributions

## Acknowledgments

This work was supported by the Army Research Office via grant W911NF-14-1-0472 and the NSF through grant BCS-1348123. The authors also thank professors T. Minor and M. Wechselberger for insightful discussions.

## Tables

**Table S1.** Dimensionless parameter values of our full model. Analogous parameters from the literature are referenced.

| Parameter | Value | Source and Ref. | Description |
| --- | --- | --- | --- |
| $n$ | 5 | assumed | Hill coefficient in upregulation function $g_c(c)$ |
| $\bar{c}_\infty$ | 0.2 | estimated from [22] | baseline stored CRH level |
| $b$ | 0.56 | estimated from [22] | relates cortisol to stored CRH level |
| $k$ | undetermined | . | relates stored CRH to CRH release rate |
| $\mu_c$ | undetermined | . | basal CRH release rate |
| $q_0$ | undetermined | . | maximum CRH release rate |
| $q_1^{-1}$ | undetermined | . | circulating CRH for half-maximum self-upregulation |
| $q_2$ | 1.8 | estimated from [21] | ratio of CRH and cortisol decay rates |
| $p_2^{-1}$ | 0.067 | $p_2^{-1}$ [43] | (*or*)-complex level for half-maximum feedback |
| $p_3$ | 7.2 | $p_3$ [43] | ratio of ACTH and cortisol decay rates |
| $p_4$ | 0.05 | $p_4$ [43] | (*or*)-complex level for half-maximum upregulation |
| $p_5$ | 0.11 | $p_5$ [43] | basal GR production rate by pituitary |
| $p_6$ | 2.9 | $p_6$ [43] | ratio of GR and cortisol decay rates |
| $t_c$ | 69.3 | assumed | CRH biosynthesis timescale |
| $t_d$ | 1.44 | "$\tau$" [43] | delay in ACTH-activated cortisol release |

## Additional Files Nondimensionalization

Our equations are nondimensionalized in a manner similar to that used by Walker *et al*. [13]:

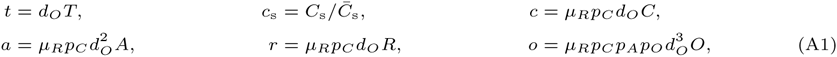

Here, *c*_*s*_, *c*, *a*, *r*, *o* are the dimensionless versions of the original concentrations *C*_*s*_, *c*, *A*, *R*, *o*, respectively. *c*_*s*_ is normalized by *C̄*_s_, which denotes the typical maximum amount of releasable CRH in the physiological range. Upon using these variables, the dimensionless forms of εEqs. 9-13 are expressed in Eqs. 14-18. The parameters *q*_*i*_,*p*_*i*_ are dimensionless combinations conveniently defined to be analogous to those used by Walker *et al*. [13]:

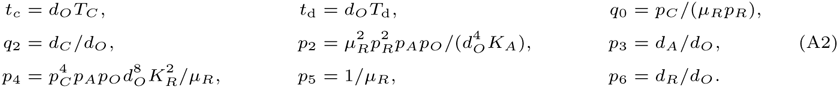

Using these scalings, we arrive at the dimensionless Eqs. 14-19.

## Parameter estimates

Many of the numerous physiological parameters in our model can be estimated or constructed from previous studies on the HPA axis. For example, as shown in Fig. A1, the parameters forming the function *c*_∞_(*o*) are derived from fitting to data on adrenalectomized male rats [22]. From the fitting, we estimate the baseline level *C̄* ≃ 0.2, and the decay rate *b* ≃ 0.6 [22]. Furthermore, the dimensionless parameters *p*_2_,…,*p*_5_ and *t*_d_ will be fixed to those used in Walker *et al*. [13]: *p*_2_ = 15, *p*_3_ = 7.2, *p*_4_ = 0.05, *p*_5_ = 0.11, *p*_6_ = 2.9 and *t*_d_ = 1.44 (*t*_d_ = 15 min). Although it is not possible to determine all of the remaining parameters from dat*a*, we will use reasonable estimates. The half-life of cortisol was estimated to be about 7.2min [21] while the half-life of CRH has been estimated to be about 4min [62]. Therefore, 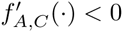. Of the remaining parameters (*n*,*μ*_*c*_, *q*_0_, *q*_1_, *k*), the dependence on n will turn out to be quantitative so we henceforth set *n* = 5. These estimated parameters are listed in Table S1. Even though one expects the values of these effective parameters to be highly variable, we fix them in order to concretely investigate the mathematical structure and qualitative predictions of our model. The parameters *μ*_*c*_, *q*_0_, *q*_1_, and *k* remain undetermined; however, it is instructive to treat *k* as a control parameter and explore the nullcline structure in *μ*_*c*_, *q*_0_, *q*_1_ parameter space.

**Figure A1:**
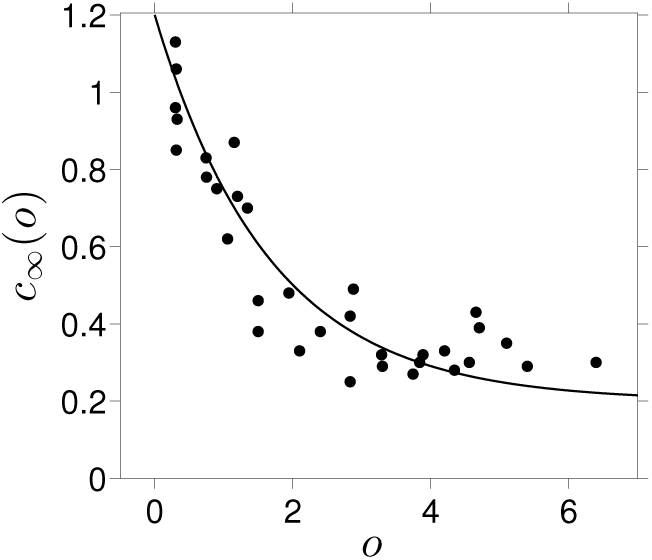
Fitting *c*_∞_(*O*) to rat data. Nondimensionalized data taken from Watts [22] and fitted using the form for *c*_∞_(*O*) given in Eq. 19. Cortisol levels were arbitrarily rescaled according to 125ng/ml = 3.

## Parameter space and nullcline structure

To determine how the *q*-nullcline crosses the *c*-nullcline, we substitute *c*_s_ by its equilibrium period-averaged value 〈*c*_∞_(*c*)〉. If we assume a basal input level *I* = 1, the values of *k* that will position the basal *q*-nullcline to just pass through the left and right bifurcation points (*q*_L_, *c*_L_) and (*q*_R_, *c*_R_) can be found by solving *q*_L,R_ = *q*_0_(1 – *e*^‒*k*〈*c*∞(*c*_L,R_))〉^):

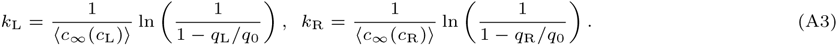

All possible ways in which the nullclines can cross each other as *k* is varied are illustrated in Fig. S2.

**Figure S2:**
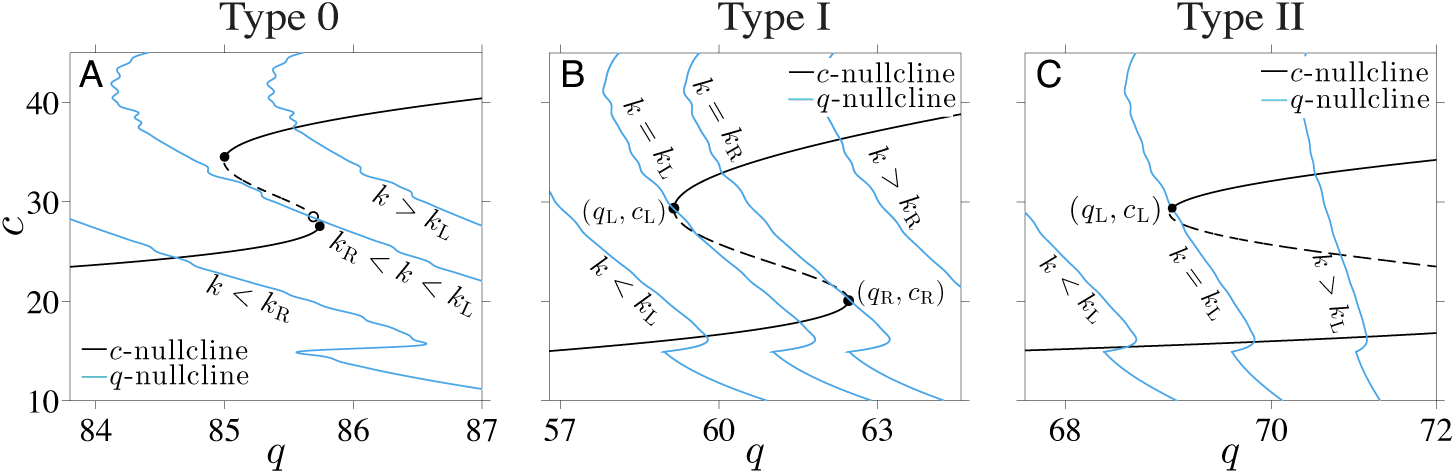
The possible number of equilibria of the reduced (*c*_*s*_, *c*) system. (A) A **Type 0** scenario in which *k*_*R*_ < *k*_*L*_ permits only one nullcline intersection, either one the lower stable branch, the unstable branch, or the upper stable branch. (B) In this **Type I** parameter regime, the *c*-nullcline is shaped and positioned such that *k*_*L*_ < *k*_*R*_. Therefore, it is possible for the model to exhibit two oscillating stable states provided *k*_*L*_ < *k* < *k*_*R*_. For *k* < *k*_*L*_ (*k* > *k*_*R*_), the *q*-nullcline shifts to the left(right) and the intersection with the upper(lower) branch of the *c*-nullcline disappears, leading to only one stable point. (*C*) A **Type II** *c*-nullcline. For *k* < *k*_*L*_, there is only one intersection at the lower branch. For all *k* > *k*_*L*_ there are two intersections.

The specific locations of the bifurcation points, as well as *k*_*L*_ and *k*_*R*_, are complicated functions of all parameters. However, Eqs. A3 allows us to distinguish three qualitatively different regimes. The first possibility is *k*_*L*_ > *k*_*R*_, where there can be at most only one intersection between the slow and fast nullclines. We denote this as a **Type 0** scenario (Fig. S2A) characterized by having at most a single stable state towards which the system will always return upon cessation of external stress. In **Type 0** situations with intermediate values of *k*, the intersection will arise in the unstable branch of the *c*-nullcline. In this case, we expect the system to oscillate between the two stable branches of the *c*-nullcline. Here, the fast variables *a*, *o*, and *r* will cycle periodically between two oscillating levels. In order for the two nullclines to intersect three times (twice on stable branches of the *c*-nullcline), the *q*-nullcline must “fit” within the bistable region of the *c*-nullcline. As shown in Fig. S2, there are two separate subcases of nullclines that intersect twice. If *k*_*L*_ < *k*_*R*_, a value of *k*_*L*_ < *k* < *k*_*R*_ would imply that the *q*-nullcline can intersect both stable branches of the *c*-nullcline, leading to two stable solutions. We refer to this case as **Type I** (Fig. S2B). Another possibility is that the right bifurcation point is beyond the maximum value *q* = *q*_0_ dictated by the function *h*(〈*c*_∞_(*c*_*R*_)〉). As shown in Fig. S2C, the bistable *c*-nullclines exhibits only one bifurcation point within the domain of *q*. The lower branch of the *c*-nullclines in this set extends across the entire range of physiological values of *q*, ensuring that the *q*-nullcline will intersect with the lower branch for any value of *k*. Therefore, to determine if there are two intersections we only need to check that *k*_*L*_ ≤ *k* is satisfied. In this **Type II** case, the system is either perpetually in the diseased low cortisol state, or is bistable between the diseased and normal states; the system will always be at least susceptible to low-cortisol disease. Summarizing,

- **Type 0**: Exactly onesolution (onenullcline intersection) existsfor the reduced subsystem. Here, *k*_*R*_ < *k*_*L*_ and the intersection may occur on the lower or upper stable branches, or on the unstable branch of the *c*-nullcline. The system is either permanently diseased, permanently resistant, or oscillates between normal and diseased states.
- **Type I**: At least one solution exists. A stable diseased solution exists if *k* < *k*_*L*_, two stable solutions (diseased and normal)arise if *k*_*L*_ ≤ *k* ≤ *k*_*R*_, and fully resistantstate arises if *k* > *k*_*R*_.
- **Type II**: At least one solution exists. A stable diseased state arises if *k* < *k*_*L*_ while both diseased and normal solutions arise if *k* > *k*_*L*_. A fully disease-resistant state cannot arise.

With the parameters fixed according to Table S1, we will treat *k* as a control parameter and exhaustively sweep the three-dimensional parameter space (*q*_0_, *q*_1_, *μ*_*c*_) to determine the regions which lead to each of the nullcline structural types. In addition, we restrict the parameter domain to regions which admit oscillating solutions of the full problem. In other words, parts of both stable branches of the *c*-nullclines must fall within values of c which support oscillations in the PA-subsystem (Fig. 3.). The regions in (*μ*_*c*_, *q*_0_, *q*_1_) space that satisfy these conditions and that yield each of the types of nullcline crossings are indicated in Fig. S3.

Based on measurements of self-upregulation of CRH secretion during stress [23], *μ*_*c*_ = 0.6 is chosen to set the baseline level of the Hill function *g*_*c*_(*c* = 0) ≈ 0.4. *q*_1_ is approximated by setting the inflection point of *g*_*c*_(*c*) to arise at *c* ≈ 25, the average value used by Walker *et al*. [13]. Assuming *c* ≈ 25 is a fixed point of Eq. 15 when *I* = 1 and *c*_*s*_ ≈ (*c*_*∞*_(25)), *q*_0_ can be estimated as a root of the right-hand-side of Eq. 15. This choice for the remaining parameters puts our nullcline system into the **Type I** category that can exhibit one or two stable states with oscillating (*a*, *o*, *r*) subsystems. We restricted the analysis of our model to **Type I** systems.

**Figure S3:**
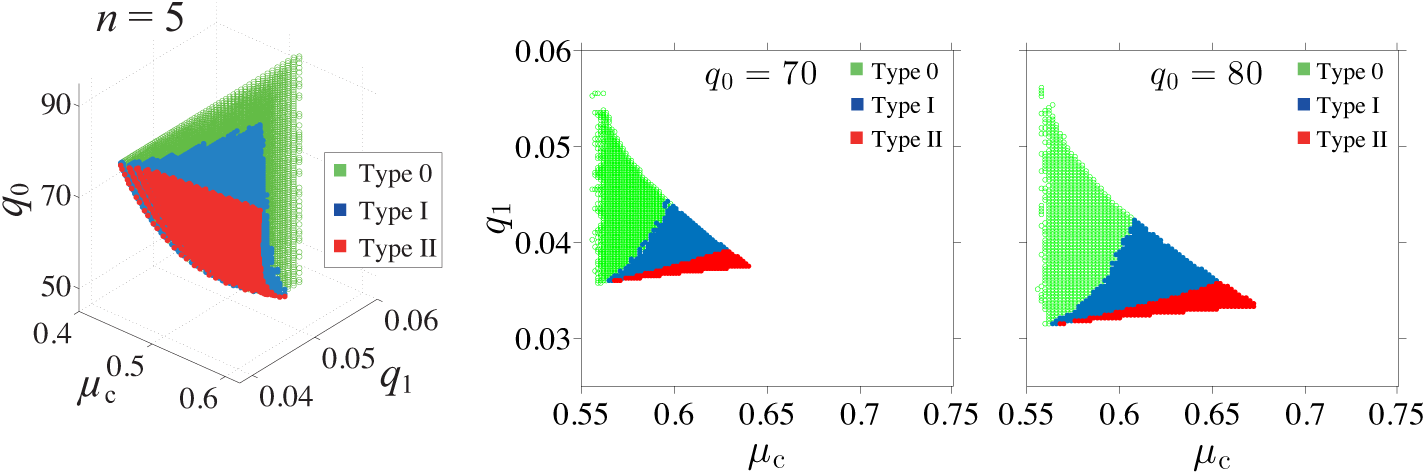
Phase diagram in (*μ*_*c*_, *q*_0_, *q*_1_)-space. Regimes for each of the three types of bistable *c*-nullclines shown in the parameter space (*μ*_*c*_, *q*_0_, *q*_1_) and (*μ*_*c*_, *q*_1_) with *n* = 5. The uncolored regions correspond to systems that do not exhibit either bistability or oscillations.

## Minimum duration and magnitude of stress

We plot the minimum duration required for normal-to-diseased transition against stress magnitude (Fig. S4). Higher magnitude of *I*_ext_ generally requires a shorter duration of stress, as expected. Note that the minimum duration is also dependent on the phase of intrinsic oscillations of the system at stress onset.

**Figure S4:**
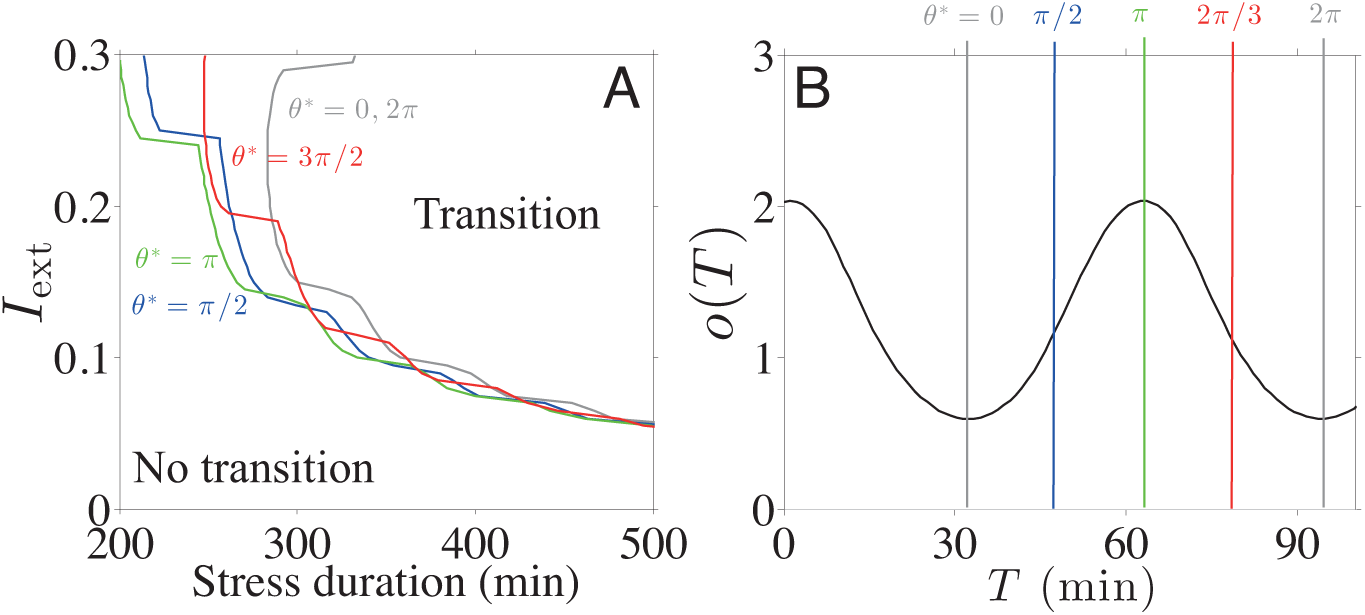
Phase diagram of stress-induced transitions. (A) Minimum duration of stress required for normal-to-disease transition is plotted against stress magnitude. The phase of intrinsic oscillations at stress onset is denoted as *θ*\*. (B) Four *θ*\* values were chosen and marked on the plot of *o*(*T*) with different colors assigned. The color of each curve in plot (A) corresponds to the *θ*^*^ of stressor onset shown in (B).

## Timing of stress onset and transient response

Here, we show how the dynamics of the system changes after the onset and cessation of stress. In previous studies [18, 21], changes in corticosterone levels in rats were measured in response to stress induced by noise applied at different phases of the animals oscillating cortisol cycle. It was observed that the timing of the stress onset relative to the ultradian phase was crucial in determining the magnitude of corticosterone response. Increases in corticosterone levels were markedly higher when noise was initiated during the rising phase than when initiated during the falling phase.

We can frame these experimental observations mechanistically within our theory. Following the experimental protocol [18, 21], we simulate the stress response using a brief stressor with a duration of 30min. As shown in Fig. S5A, an external stress that is applied mostly over the falling phase of the cortisol oscillation results in a higher subsequent nadir in *o*(*t*) than one that is applied predominantly during a rising phase. However, as shown in Fig. S5B, stress applied mainly during the rising phase leads to a higher subsequent peak level. This observation is consistent with the results of the experiment on rats and can be explained by the dynamics inherent in our model. The immediate increase in *q* = *q*_*o*_/*h*(*c*_*s*_) associated with the increase in *I* leads to a rapid increase in *c*, as shown in Fig. 7.. This higher level of circulating CRH shifts the stable limit cycle of the PA subsystem to a new one with higher minimum and maximum values of ACTH and cortisol (as shown in Fig. 3.). This new limit cycle is shown by the blue curve in Figs. S5C,D. Under external stress, a trajectory of the system quickly deviates and approaches the new limit cycle, but quickly returns to the original limit cycle after cessation of stress. Thus, depending on the position of the trajectory relative to that of the new stressed limit cycle, the initial deviation may try to reach the new limit cycle in the falling or rising cortisol phases as shown in Figs. S5C,D. Moreover, if the duration of the stress is shorter than the period of the inherent oscillation, the trajectory will return to its original limit cycle before completing a full cycle of the new limit cycle. These properties of the limit cycle dynamics explain the difference in the level of subsequent peak following the stress onset depending on the timing of the stress onset.

**Figure S5:**
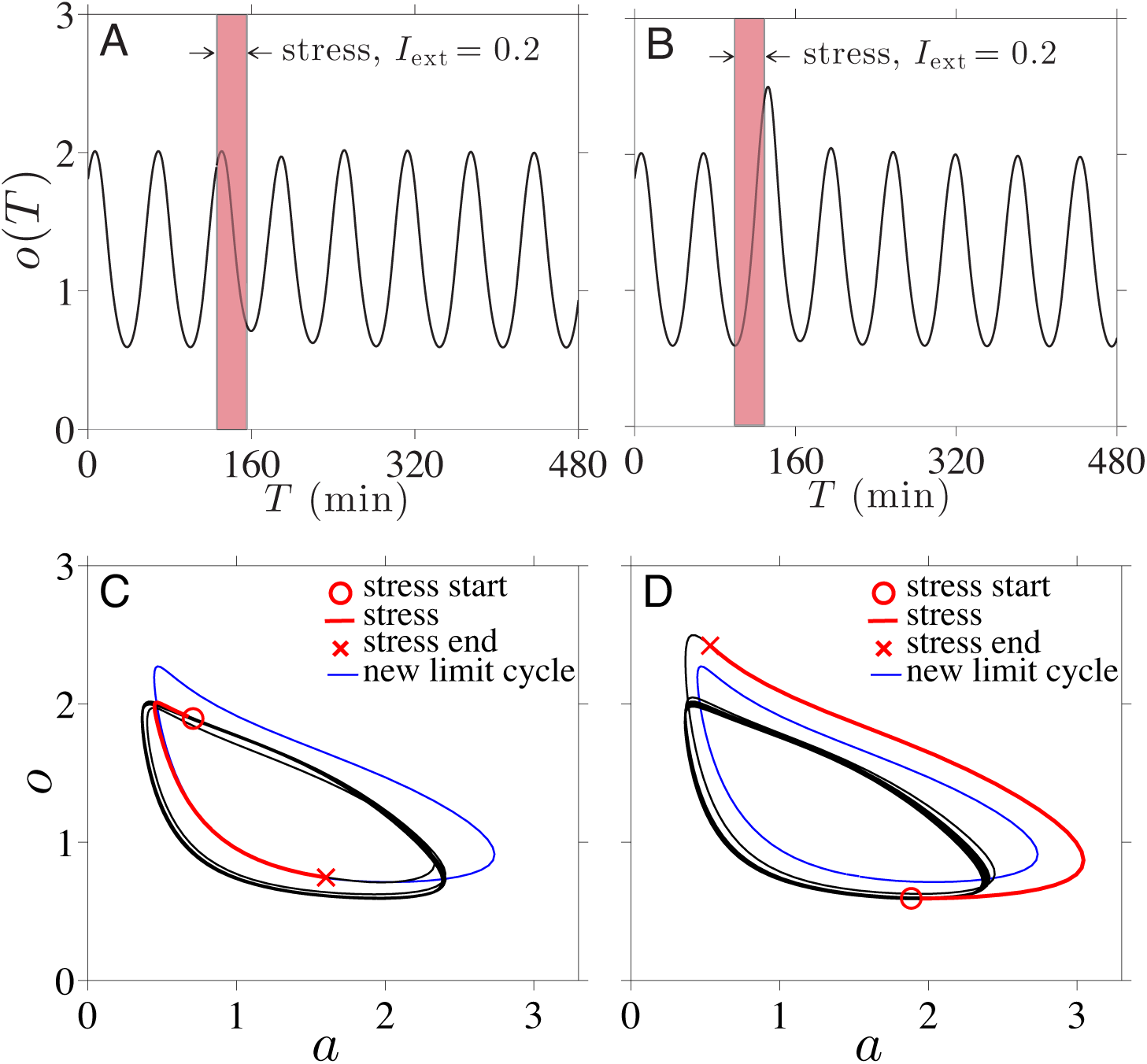
Stress timing and cortisol response. (A) A stressor of duration of 30min with magnitude *I*_ext_ = 0.2 was applied mainly over the falling phase of the underlying cortisol oscillation. The first peak after the stress onset was almost unchanged, but the first nadir was elevated. (B) The same stressor used in (A) applied during the rising phase led to a significantly increased subsequent peak while the first nadir was unaffected. (*C*) The trajectory of the system (red) is projected onto the cortisol-ACTH plane. The new limit cycle of the PA-subsystem corresponding to fixed *I*(*t*) = 1.2 is indicated by the blue curve. During stress, the trajectory of the system is attracted towards the new limit cycle. The system recovers after making a smaller cycle within the normal limit cycle, reaching a higher nadir. (D) The trajectory of the system deviates then recovers back through a trajectory above the normal limit cycle, reaching a higher peak.

## Cortisol dependent Iext

As it has been shown that synaptic input of the PVN cells is modulated by cortisol for certain types of stressor, we briefly discuss how cortisol dependent *I*_ext_(*T*, *O*) may affect the behavior of our model. Within our model, modulation in synaptic input by glucocorticoids can be viewed as a cortisol dependent external input function: *I*^ext^ (*T*, *O*) = /time (*T*) + /cort(*O*). One possible form of /ext (*T*, *O*) is illustrated in Fig. S6 where *I*_ext_(*T*, *O*) is assumed to be lower when cortisol levels are higher. Since it was shown that cortisol does not affect the basal release rate [37], the cortisol dependent component of the external input function, *I*_cort_(*O*), should be zero when there is no stress. On the other hand, it was also shown that the inhibition effect cannot decrease the release rate below the basal rate [37] so we can further assume that *I*_ext_(*O*, *T*) ≥ 0. When these conditions are met, the modification in *I*_ext_(*T*, *O*) should not affect the bistability of the system since *I*(*T*) = *I*_base_ = 1 is unchanged. However, a cortisol dependent *I*_ext_(*T*, *O*) will make the timing of stress onset become more relevant in predicting whether or not stressors can induce transitions between normal and diseased states. Driven by the intrinsic oscillations in *O*(*T*), *I*_ext_(*T*) will also oscillate during stress, leading the *q*-nullcline to shift back and forth during stress in the (*q*, *c*)-plane as shown in Fig. S6C. Oscillations in the *q*-nullcline affect the net decrease in *q* during stress, changing the position of the system on the (*q*, *c*)-plane relative to the separatrix between the normal and the diseased basins of attraction at stress termination.

**Figure S6:**
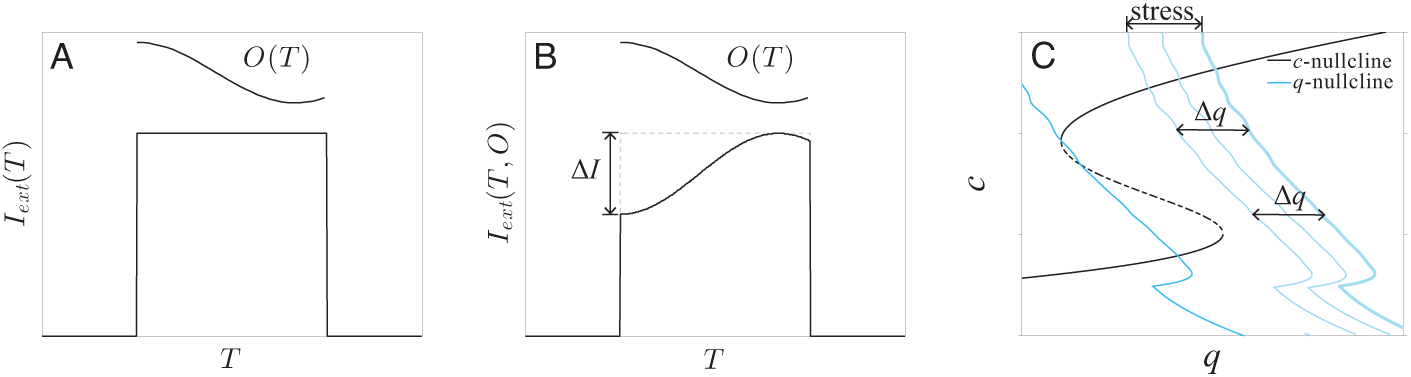
Cortisol dependent synaptic input of the PVN and its possible effects. (A) Cortisol independent *I*_ext_(*T*) used in our current model. (B) An example of cortisol dependent *I*_ext_(*T*, *O*), where we assume the synaptic input of the PVN is attenuated at higher levels of *O*(*T*). (*C*) During stress, the *q*-nullcline shifts back and forth in the (*q*,*c*)-plane due to oscillations in *I*_ext_(*T*, *O*) as driven by the intrinsic ultradian oscillations in *O*(*T*). In turn, theses shifts in *q*-nullcline will affect the net decrease in *q* during stress. Since transitions are sensitive to the position of *q* at stress termination, including a cortisol dependent *I*_ext_(*T*, *O*) will make transitions more strongly dependent on the timing of stress onset.

